# Single cell profiling reveals novel tumor and myeloid subpopulations in small cell lung cancer

**DOI:** 10.1101/2020.12.01.406363

**Authors:** Joseph M Chan, Álvaro Quintanal-Villalonga, Vianne Gao, Yubin Xie, Viola Allaj, Ojasvi Chaudhary, Ignas Masilionis, Jacklynn Egger, Andrew Chow, Thomas Walle, Marissa Mattar, Dig VK Yarlagadda, James L. Wang, Fathema Uddin, Michael Offin, Metamia Ciampricotti, Umesh K Bhanot, W Victoria Lai, Matthew J Bott, David R Jones, Arvin Ruiz, Travis Hollmann, John T Poirier, Tal Nawy, Linas Mazutis, Triparna Sen, Dana Pe’er, Charles M Rudin

## Abstract

Small cell lung cancer (SCLC) is an aggressive malignancy that includes subtypes defined by differential expression of *ASCL1*, *NEUROD1*, and *POU2F3* (SCLC-A, -N, and -P, respectively), which are associated with distinct therapeutic vulnerabilities. To define the heterogeneity of tumors and their associated microenvironments across subtypes, we sequenced 54,523 cellular transcriptomes from 21 human biospecimens. Our single-cell SCLC atlas reveals tumor diversity exceeding lung adenocarcinoma, driven by canonical, intermediate, and admixed subtypes. We discovered a *PLCG2*-high tumor cell population with stem-like, pro-metastatic features that recurs across subtypes and predicts worse overall survival, and manipulation of *PLCG2* expression in cells confirms correlation with key metastatic markers. Treatment and subtype are associated with substantial phenotypic changes in the SCLC immune microenvironment, with greater T-cell dysfunction in SCLC-N than SCLC-A. Moreover, the recurrent, *PLCG2-*high subclone is associated with exhausted CD8+ T-cells and a pro-fibrotic, immunosuppressive monocyte/macrophage population, suggesting possible tumor-immune coordination to promote metastasis.

## INTRODUCTION

Small cell lung cancer (SCLC) is the most aggressive lung cancer subtype, responsible for approximately 25,000 deaths annually in the US and an estimated quarter million deaths worldwide^1,2^. The prognosis for SCLC patients remains exceptionally poor: most patients present with metastatic disease, and the recent addition of immune checkpoint blockade to first-line chemotherapy has only modestly improved median survival to roughly one year^3,4^. The strong predilection for early metastasis and the rapid emergence of therapeutic resistance contribute to poor long-term outcomes, with 5-year survival of 15-30% for limited stage disease, and less than 1% for extensive disease^1,2^.

Although SCLC appears morphologically homogeneous, recent data from both murine models and human tumors suggest the existence of SCLC subtypes with distinct therapeutic vulnerabilities^5^. These subtypes have been classified based on differential expression of four transcription factors (TFs): *ASCL1*, *NEUROD1*, *POU2F3* or *YAP1*^5^. The emerging consensus on SCLC subtypes^5^ has led to new questions, such as whether subtypes are associated with different disease stages, metastatic potential or immune microenvironments, and whether there is plasticity between subtypes^5–7^.

Single cell RNA sequencing (scRNAseq) offers a unique opportunity to address these questions by dissecting the intratumoral heterogeneity and surrounding tumor microenvironment (TME) of SCLC. Efforts to apply this technology to human SCLC tumors have been limited, as surgical resections of primary tumors are performed in under 5% of SCLC patients^8^, and biopsied samples are not typically preserved in a manner amenable to single-cell profiling. Since resection is only clinically indicated for early stage *de novo* disease, these samples fail to capture the spectrum of disease progression.

To address these limitations, we have optimized the processing of thoracenteses, core needle biopsies, and fine needle aspirates, and applied these approaches to construct the first single-cell atlas of SCLC patient biospecimens, with comparative lung adenocarcinoma (LUAD) and normal lung data. Our analysis reveals unexpectedly high inter-patient transcriptomic heterogeneity of cellular states in SCLC tumor and immune cells. We delineate subtype-specific gene programs, describe prevalent promiscuity between subtypes, and find a particularly immunosuppressed TME in SCLC compared to LUAD, with additional subtype-specific changes in immune dysfunction. In the midst of substantial heterogeneity, we identify a remarkable stem-like pro-metastatic tumor subpopulation marked by high *PLCG2* expression that spans the full diversity of SCLC subtypes and predicts worse overall survival. Together, our analyses provide a deep characterization of the molecular features of SCLC, with clinical implications.

## RESULTS

### The human SCLC single-cell transcriptional landscape reveals unexpected tumor heterogeneity

We profiled the transcriptomes of 155,098 cells from 21 fresh SCLC clinical samples obtained from 19 patients, as well as 24 LUAD and 4 tumor-adjacent normal lung samples as controls (**Figures 1A** and **S1A**). The SCLC and LUAD cohorts (**Table S1**) include treated and untreated patients (**Figure 1B**). Samples were obtained from primary tumors, regional lymph node metastases, and distant metastases (liver, adrenal gland, axilla, and pleural effusion) (**Figure 1C**).

**Figure 1:**
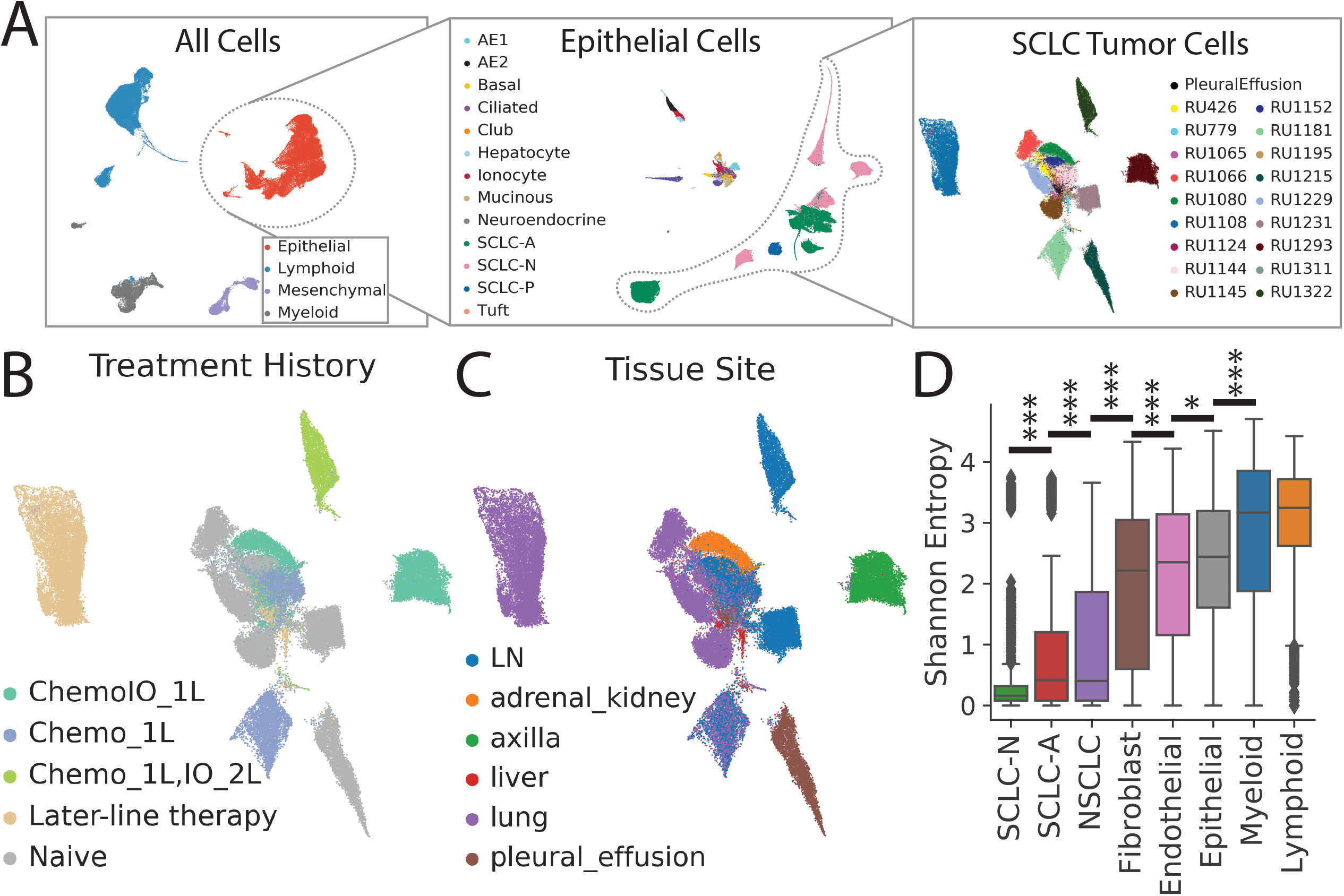
The single-cell transcriptional landscape of SCLC. Related to Supplementary Figure S1 and Supplementary Table S1. LUAD and normal adjacent lung serve as reference tissues. (A) UMAP projections of iterative subsets of cells from the global level (left, n=155,098 cells) to the epithelial compartment (middle, n=64,301 cells) to SCLC tumor cells (right, n=54,523 cells). Each dot represents a single cell colored by cell type. (B) UMAP projection of SCLC tumor cells annotated by treatment history. (C) UMAP projection of SCLC tumor cells annotated by tissue site. (D) Inter-patient heterogeneity, as measured by the Shannon entropy of samples for each cell type, computed by bootstrapping to correct for the number of cells in each cluster and cell type (**STAR Methods**). Cell types are ordered by median Shannon diversity and significant differences in entropy between neighboring cell types are shown (Mann-Whitney test). DC = dendritic cells, LN = lymph node, Chemo_1L = chemotherapy in first line, ChemoIO_1L = chemotherapy plus immunotherapy in first line, IO_2L = Immunotherapy in second line, later-line therapy = multiple lines of treatment. p-values: *<0.05, **<0.01, ***<0.001.

All scRNA-seq data were merged, normalized, batch-corrected, and clustered to identify coarse cell types, including epithelial, mesenchymal, lymphoid, and myeloid cells (**Figure 1A, S1A; STAR Methods**). Further clustering within the epithelial compartment identified cells comprising the respiratory epithelium (including alveolar epithelial types 1 and 2, ciliated, club, neuroendocrine and tuft cells) and hepatocytes derived from our liver metastases (**Figures 1A and S1A; STAR Methods**).

MSK-IMPACT™ targeted sequencing^9^ of 13 SCLC samples shows frequent mutation or loss of *RB1* and *TP53*, and recurrent mutations in *CREBBP* and *KMT2B* (**Figure S1B and S1C**). This information facilitated the identification of tumor cells that harbor reads bearing the genomic variant. We also inferred single-cell copy number to support tumor cell identification **(STAR Methods)**. Inferred copy number variation (CNV) levels are higher in SCLCs than LUAD (**Figures S1D and S1E)**, consistent with SCLC having a higher tumor mutation burden^10^. Based on previous studies investigating the most likely cell types of origin^11^, we consider neuroendocrine and alveolar epithelial type 2-like clusters to represent SCLC and LUAD tumor cells, respectively.

Following cell type annotation, we characterized the tumor heterogeneity within our atlas. Of 38 epithelial clusters (64,301 cells), we found that LUAD and SCLC clustered separately as expected, with 5 LUAD clusters comprised of 7,635 cells from 24 tumors and 25 SCLC clusters comprised of 55,815 cells from 21 tumors, consistent with lower tumor purity in LUAD (**Figure S1F**). To quantify the inter-patient heterogeneity of SCLC, we calculated the Shannon diversity of patients per cluster and compared the diversity of SCLC tumor cells to LUAD and normal stroma. For a given cluster, a low Shannon diversity signifies that the phenotype is rarely shared across patients, indicating high inter-patient heterogeneity. SCLC tumors showed significantly lower Shannon diversity compared to LUAD (**Figure 1B**). We also observed high Shannon diversity in stromal and immune cell populations, consistent with minimal batch effects across samples, and low Shannon diversity (high phenotypic diversity) in tumor cells compared to stroma, consistent with prior studies^12,13^. Our results suggest that, despite its homogeneous histological morphology, SCLC has a high degree of transcriptional tumor heterogeneity, exceeding that of LUAD and normal stroma.

### Tumor heterogeneity of canonical and non-canonical SCLC subtypes at single-cell resolution

Next, we characterized cell states within the major SCLC subtypes^14^, including only the SCLC tumor compartment (54,523 cells) and restricting our feature selection to gene signatures of canonical SCLC subtypes that we identified from an independent bulk RNA-seq cohort (**Tables S2, S3, S4 and S5; STAR Methods**).

SCLC subtypes are typically classified by the expression of canonical TFs (*ASCL1*, *NEUROD1*, *POU2F3*, *YAP1*), but a strategy dependent on a single gene is unreliable given the gene dropout prevalent in scRNA-seq. We therefore used a neighbor-graph-based approach to calculate the probability of a given SCLC subtype per cell^15^ (**STAR Methods**), which can harness the full complexity of each subtype composed of multiple gene programs. We identified the most likely subtype of each cell (**Figure 2A**) and used this information to categorize the major subclone of each sample as subtype SCLC-A (N = 14), SCLC-N (N = 6), or SCLC-P (N = 1) (**Figure 2A, STAR Methods**). We did not observe any YAP1-expressing tumor cells, consistent with absence of SCLC-Y. Our classification is supported by the relative expression of canonical TFs (**Figure 2B**) and matched immunohistochemistry (IHC) when available (**Figure S1G)**. However, unlike single-gene expression or IHC, our strategy is also able to classify “double-positive” cases (*ASCL1-*high *NEUROD1*-high Ru1231 as SCLC-N) and “double-negative” cases (*ASCL1-*low *NEUROD1-*low Ru1293A as SCLC-N; notably, Ru1293A retains *NEUROD2* and *NEUROD4* expression).

**Figure 2:**
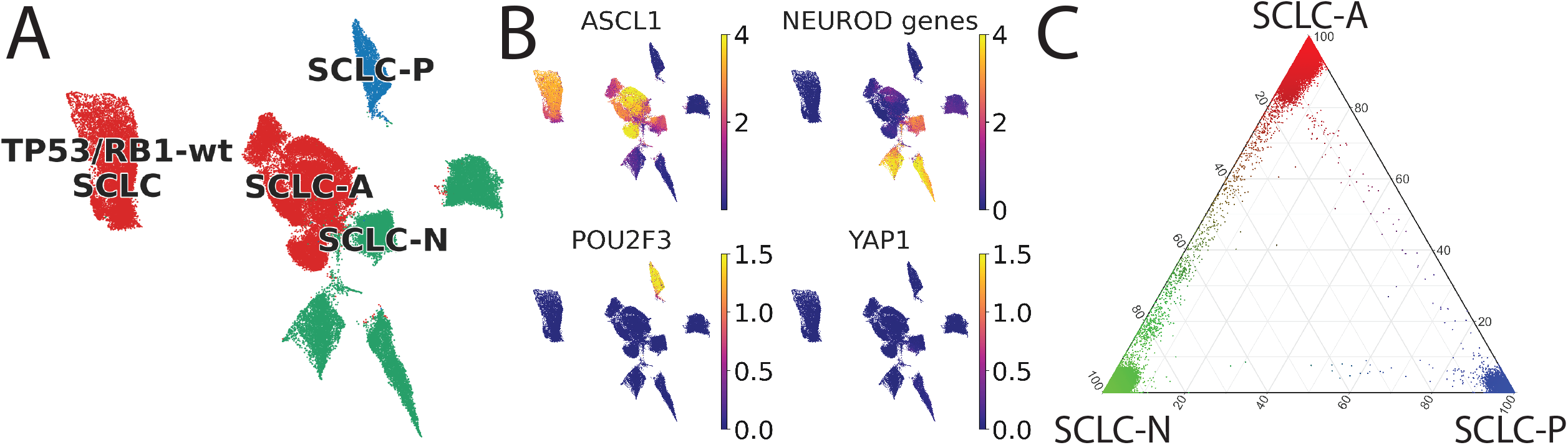
Single-cell sequencing resolves SCLC subtypes in patient samples at high resolution. Related to Supplementary Figures S1-2 and Supplementary Table S1. (A) UMAP projection of SCLC cells colored by subtype (red = SCLC-A, green = SCLC-N, blue = SCLC-P), based on maximum likelihood computed by our classifier (**STAR Methods**). Sample RU1108a is labeled as a *TP53/RB1* wild-type SCLC-A outlier. (B) UMAP projection of MAGIC-imputed^108^ expression of *ASCL1*, *NEUROD1*, *POU2F3* and *YAP1* in the SCLC cohort (**STAR Methods**). Expression in units of log2(X+1) where X = normalized counts. (C) Ternary plot of SCLC subtype probability per cell, calculated by Markov absorption probabilities (**STAR Methods**). Cell color is assigned by the likelihood of SCLC-A (red), SCLC-N (green), and SCLC-P (blue).

The vast majority of tumors are predominantly composed of a single SCLC subtype (**Figure S1H**). However, most samples also include a minority of cells that fall along a continuous spectrum from SCLC-A to SCLC-N (**Figure 2C; STAR Methods**). One sample (RU1215) harbors discrete populations of both SCLC-A and SCLC-N (**Figure S2A**). Our analysis identified cells in apparent transition between known subtypes (**STAR Methods**), which may represent non-canonical phenotypes or intermediate subtype states. These findings are consistent with our previous report of ASCL1+/NEUROD1+ cells in SCLC clinical samples^16^ and other reports showing SCLC-A to SCLC-N transitions upon disease progression in SCLC preclinical models^7,17^.

### SCLC-N exhibits a pro-metastatic neuronal and EMT phenotype

To understand the role of SCLC subtype in tumor progression, we assessed cell composition and gene expression differences across subtypes (**Figure S2B**). Consistent with observations in mouse models^7,17^, we found SCLC-A to be significantly overrepresented in primary tumors whereas non-SCLC-A subtypes are enriched in nodal and distant metastases (Dirichlet regression, p<3.4×10^−8^; **Figure S2C; STAR Methods**). We also observed greater interpatient diversity in SCLC-N tumors as compared to SCLC-A (**Figure 1D**). These findings are consistent with preclinical models showing SCLC-N can progress from SCLC-A through discrete evolutionary bottlenecks.

We performed differential expression (DE) and pathway analysis to determine subtype-specific gene programs (**Figures 3A and S2D; and Tables S6, S7, S8, S9, S10 and S11**). We found that SCLC-A is enriched in expression of genes regulating cell cycle progression and DNA repair, as well as *EZH2* target genes implicated in SCLC cell cycle^18,19^ (**Figure S2D**). In contrast, SCLC-N tumors exhibit a pro-metastatic phenotype with overexpressed markers of epithelial-mesenchymal transition (EMT) (*VIM, ZEB1* and *TWIST1*)^20^ and hypoxia and angiogenesis (*HIF1A, VEGFA* or *FOXO3*) (**Figures 3A, 3B and S2D**). SCLC-N also overexpressed metastasis-related signaling pathways, including (1) TGF-β^21^ (upregulation of *TGFB1* and *TFGBR1/3*); (2) BMP signaling^20,22^ (upregulation of ligands *BMP2/7* and receptors *BMPR1A/2*)^23^; (3) STAT signaling, (upregulation of *STAT3*, *IL6R*, *IL11RA*)^20^; and (4) TNFα-mediated NFκB signaling (upregulation of *TNF, SMAD3, PHLDA1*)^24,25^ (**Figures 3A, 3B and S2D**).

**Figure 3:**
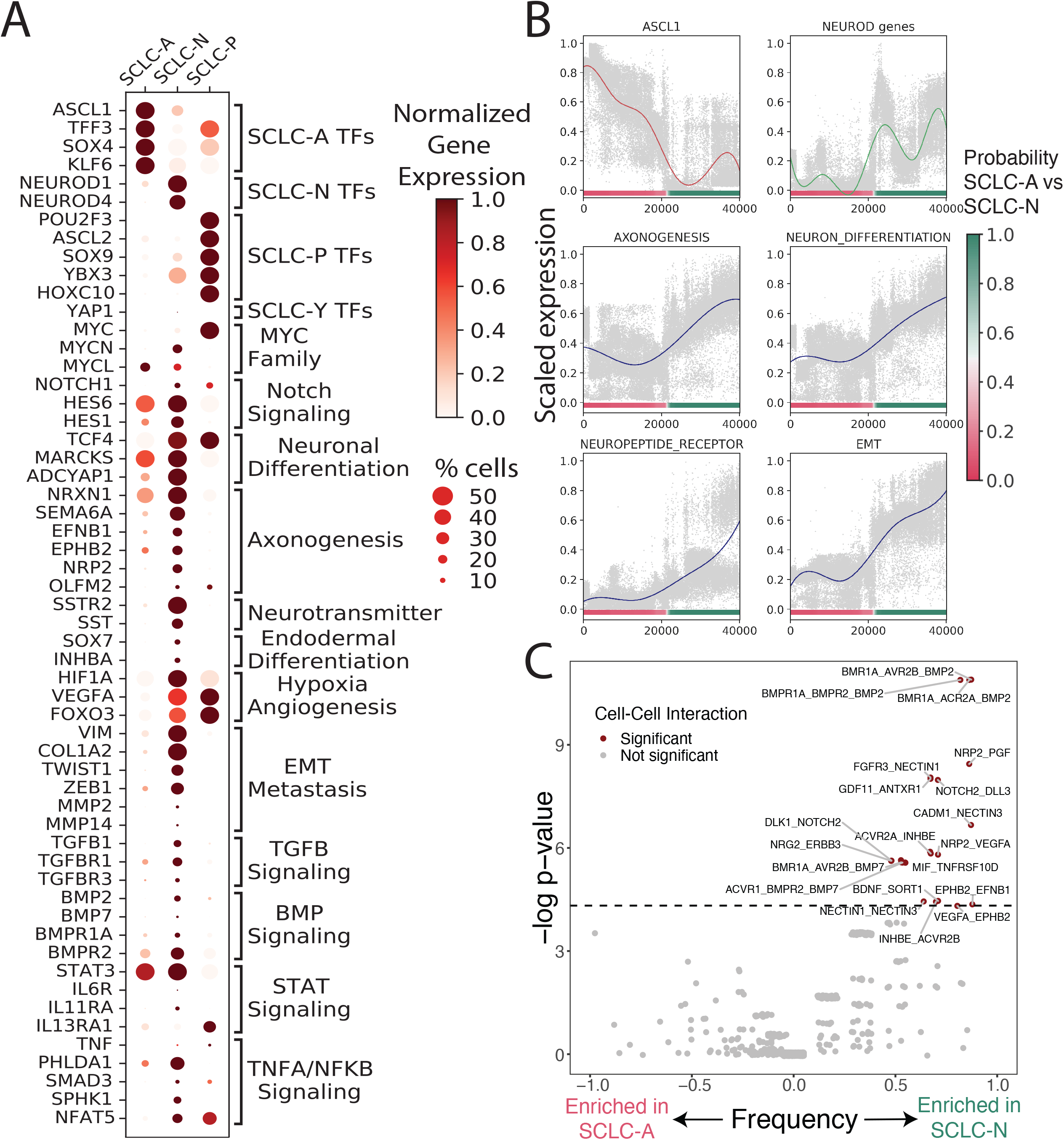
Gene programs and cell-cell interactions enriched in each SCLC subtype. Related to Supplementary Figure S2 and Supplementary Table S1. (A) Dot plot showing selected DEGs between each SCLC subtype versus the rest, as well as between SCLC-A vs SCLC-N. DEGs are grouped by enriched gene pathways as assessed by GSEA (NES > 1, FDR < 0.1) (**STAR Methods, Table S6**). Dot size = % cells expressing gene; dot color = mean expression scaled from 0 to 1. (B) Scaled expression of canonical markers, or scaled average Z-score of select enriched pathways in SCLC-N (Y-axis), versus SCLC subtype probability (X-axis). Solid lines represent average gene/pathway trend (**STAR Methods**). (C) Enrichment of tumor-tumor interactions within SCLC-A vs SCLC-N. Significant interactions are first assessed using CellPhoneDB^109^. Enrichment of interactions within SCLC-A vs SCLC-N is then plotted as significance (−log_2_ of Fisher’s test) versus frequency (**STAR Methods**). Dashed line corresponds to nominal p < 0.05.

SCLC-N displayed a neuronal differentiation phenotype, with high expression of key neurogenesis factor *TCF4*^26,27^ involved in BMP signaling and metastasis^28,29^, as well as a neuropeptide signaling signature (*SSTR2, SST* or *MARCKS*) (**Figures 3A and 3B and Table S6**). Interestingly, we found that SCLC-N was enriched in two main axonogenic signaling pathways: ephrin (*EFNB1* and *EPHB2*, among others)^30^ and semaphorin (*SEMA6A* and *NRP2*, among others)^31^. Prior studies have shown that the axonogenesis program coordinates cell polarity with neuronal migration^32^; it has been implicated in SCLC metastasis^33^, but never before in SCLC-N specifically. We have shown that LUAD hijacks endodermal developmental pathways in metastasis^34^; similarly, our findings here suggest that SCLC-N may adopt a neuronal developmental phenotype to achieve a metastatic state.

We further assessed differentially expressed ligand-receptor pairs between subtypes (**Figure 3C**; **STAR Methods**), and observed enrichment in homotypic (tumor-tumor) interactions in SCLC-N compared to SCLC-A. While one can not be certain of any individual hypothesized ligand-receptor interaction in such analysis, the difference in number of interactions between subtypes is striking. This enrichment is consistent with growth patterns observed *in vitro*; SCLC-A cell lines typically grow as loose floating aggregates, whereas SCLC-N lines more commonly grow as a tightly adherent monolayer in cell culture^35^. The biphenotypic sample, Ru1215, allowed us to explore interactions between SCLC-A and -N subtypes in the same tumor (**Figures S2A and S2E; STAR Methods**). Ligand-receptor analysis in this sample suggested multiple interactions between the two subtypes that recapitulated findings previously shown only in preclinical models^35,36^.

### A novel stem-like, pro-metastatic cell cluster recurs across patients and SCLC subtypes

The transcriptomic diversity of SCLC contrasts with the uniformly poor prognosis of patients. We analyzed phenotypes spanning multiple patients to determine whether any shared cell types may account for the universal aggressiveness of SCLC. Unsupervised clustering of the SCLC tumor compartment identified 25 clusters, most of which are specific to a single sample. However, cluster 22 is the most highly recurrent cluster across samples (Mann-Whitney p < 2.2×10^−16^), spanning a range of treatment statuses, tissue sites (**Figures 4A and 4B; Table S1; STAR Methods**), and major subclonal subtypes (**Figure 4C**). Cells in the recurrent cluster exhibit significantly higher uncertainty in subtype assignment than those in any other cluster (Mann-Whitney p < 2.2×10^−16^), suggesting a dedifferentiated phenotype (**Figure 4A; STAR Methods**). DE and pathway analysis showed that this phenotype is enriched in gene programs related to metastasis and stemness (**Figures 4D and 4E, Table S12**).

**Figure 4:**
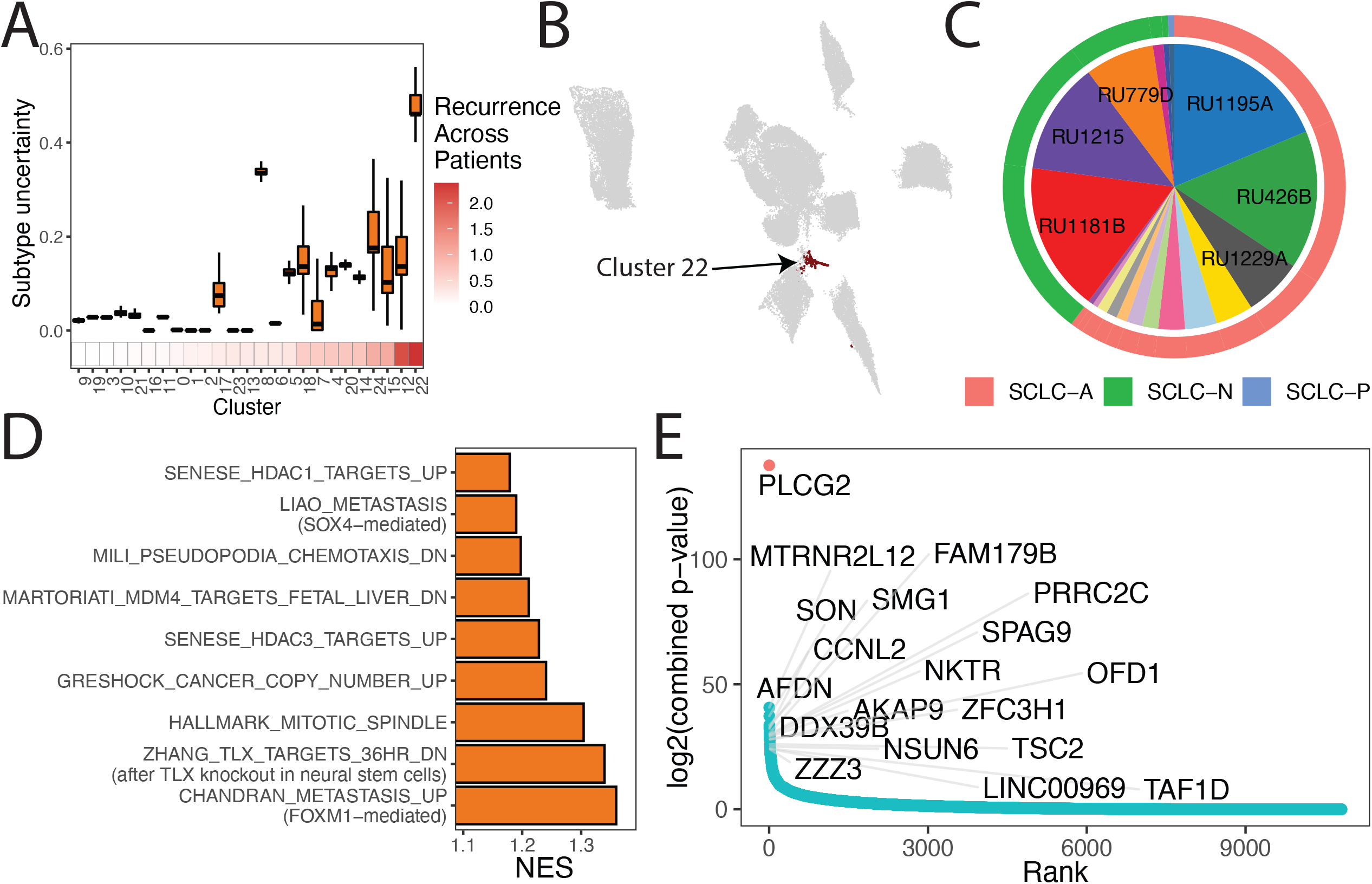
A subpopulation with metastatic, stem-like phenotype recurs broadly across SCLC tumors. Related to Supplementary Figure S3 and Supplementary Table S1. (A) Box plot of subtype uncertainty per SCLC tumor cluster, ordered by recurrence across patients. For each cell, uncertainty is scored as entropy of SCLC subtype probabilities. The distribution of cellular uncertainties in each cluster is shown. Recurrence across patients is scored as Shannon entropy of patients within the cluster (**STAR Methods**). (B) UMAP projection highlighting cells in recurrent cluster 22 (black). (C) Proportion of samples comprising the recurrent cluster (9 of 23 profiled patients harboring at least 3% of the cluster). Outer ring indicates the major subclonal subtype of a sample. (D) Gene programs significantly enriched in cluster 22. Bar plot of NES from GSEA for each pathway. Significantly enriched pathways have FDR < 0.05 and NES > 1 (**Table S12**). (E) Genes ordered from most to least recurrently overexpressed along the X-axis, with recurrence score plotted on the Y-axis. The recurrence score is calculated as follows. Within each sample, DEGs were assessed between the recurrent cluster versus the rest of the tumor. The associated adjusted p-values for differential expression within each tumor are combined using Edgington’s method. The recurrence score is the −log of the combined p-value (**Table S14**) (**STAR Methods**).

Within Cluster 22, *phospholipase C gamma 2* (*PLCG2*) is the top upregulated gene (**Figures 4E and S3A, Tables S13 and S14**). *PLCG2* has been previously implicated in Alzheimer’s disease^37,38^ and its paralog *PLCG1* promotes metastasis^39,40^. We used knnDREMI^41^, which is well suited to handle data sparsity and rare cell populations, to explore the full gene program that covaries with *PLCG2* (**STAR Methods**), and grouped results into three gene modules corresponding to low (module 1), medium (module 2) and high *PLCG2* expression (module 3) (**Figure S3B and Table S15**). Candidate genes in module 3 included *FGFR1* (implicated in SCLC through frequent amplifications^42^, and *MTRNR2L8* and *MTRNR2L12* (humanin family genes shown to inhibit apoptosis^43^, to be neuroprotective in Alzheimer’s disease^44^, and to promote tumor progression in triple-negative breast cancer^45^). Among the top 5% of pathways most correlated to module 3 were those related to stemness (including *OCT4* and *SOX2* targets), metastatic gene signatures, and pro-metastatic signaling pathways (including Wnt and BMP signaling)^20^ (**Figures S3B,C and Table S16**).

### The PLCG2+ recurrent cluster is associated with reduced overall survival in SCLC patients

While our recurrent SCLC cluster was characterized by the overexpression of multiple genes (**Figure S3A**), we investigated the role of *PLCG2* expression as a proxy of this subpopulation in our analyses, as this gene showed striking overexpression in this subpopulation. Consistent with the pro-metastatic phenotype of the shared cluster, we found that *PLCG2* alone is significantly upregulated in metastatic sites compared to lung, and rises further in the liver, the most common metastatic location for SCLC (**Figure 5A**). These observations prompted us to test *PLCG2* function directly by knocking out *PLCG2* in a high-*PLCG2* SCLC cell line (DMS114) and overexpressing *PLCG2* in an SCLC cell line with relatively low expression (H82). In a transwell assay, overexpression increases migration (p = 0.030) and invasion (p = 0.019) in H82 cells, and knockout suppresses migration (p = 0.019) and invasion (p = 0.002) in DMS114 cells (**Figures 5B and 5C; STAR Methods**). *PLCG2* expression in these cell lines is associated with higher Wnt and BMP signaling (**Figure S3D**), pro-metastatic pathways^20^ that we found are highly correlated with *PLCG2*-high cells (**Figures S3B and S3C**). In line with these results, *PLCG2* induction upregulates EMT markers in H82 (**Figure S3D**). No expression of any of the EMT markers tested was detected in DMS114 (data not shown).

**Figure 5:**
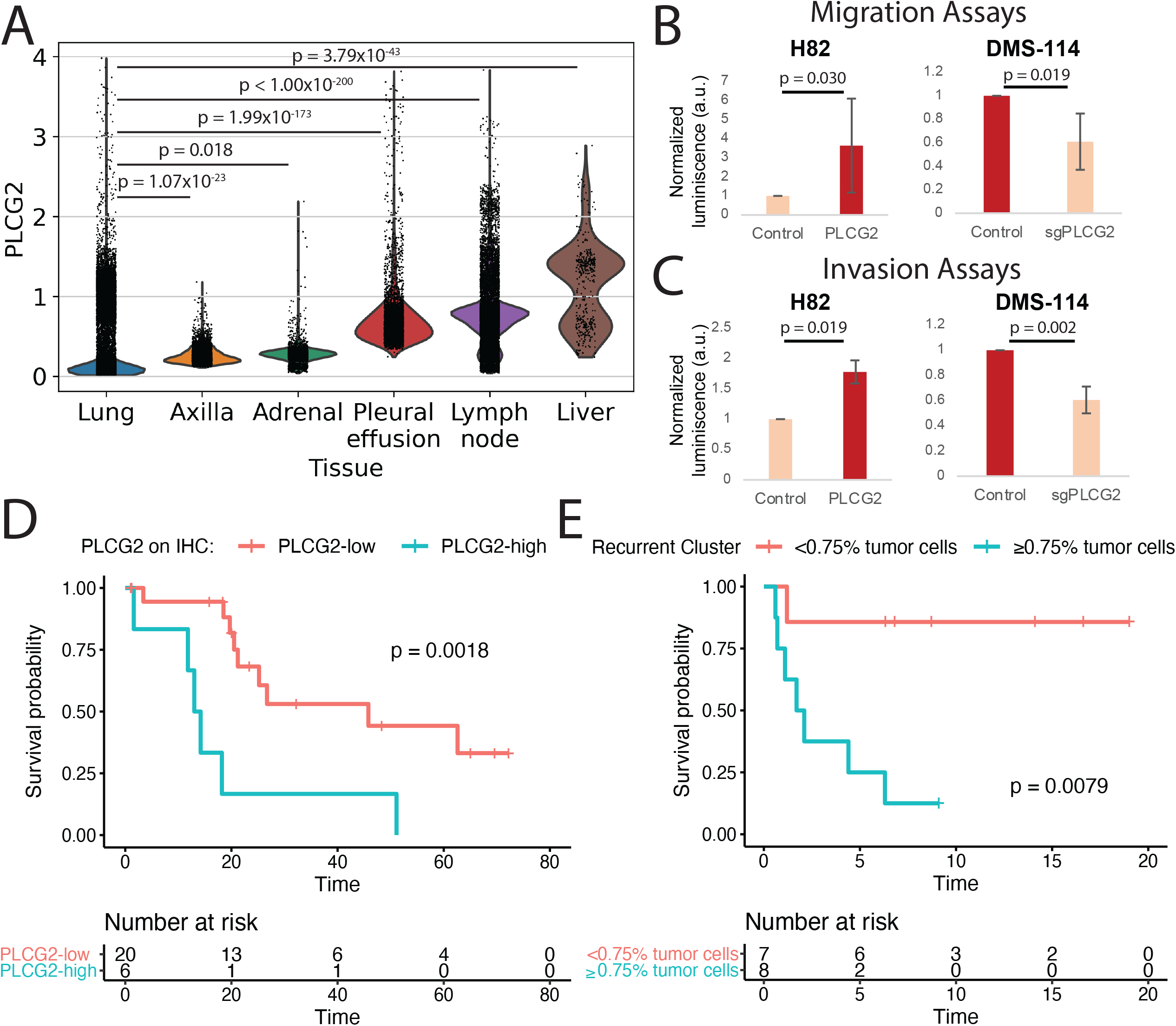
A role for the PLCG2+ recurrent cluster in metastasis and patient outcome partially mediated by PLCG2 expression. Related to Supplementary Figure S3 and Supplementary Tables S1-3. (A) Violin plot with *PLCG2* expression in our SCLC samples, grouped by tissue site. Bonferroni-adjusted p-values were calculated using Mann-Whitney test. Expression is plotted as log2(X+1) where X is the normalized count, imputed using MAGIC with k=30, t=3. (B) Migration assays for *PLCG2*-overexpressing H82 and *PLCG2*-downregulated DMS-114 cell lines. Migration capacity was measured with a luminometric method in a minimum of 3 independent experiments, and normalized to control condition, which takes the value of 1. Differences in migration capacity were assessed by two-tailed Student’s t-test. (C) Invasion assays for *PLCG2*-overexpressing H82 and *PLCG2*-downregulated DMS-114 cell lines. Invasion capacity was measured with a luminometric method in a minimum of 3 independent experiments (3 technical replicates/experiment), and normalized to control condition, which takes the value of 1. Average +/− SD values of all experiments are shown. Differences in invasion capacity were assessed by two-tailed Student’s t-test. (D) Kaplan-Meier analysis of OS in an independent cohort of SCLC patients (**Table S17**) in patients with high vs low *PLCG2* positivity (>15% vs ≤15% of tumor cells with high *PLCG2* staining intensity), as assessed by IHC performed on a TMA. (E) Kaplan-Meier analysis of overall survival (OS) in patients with detectable *PLCG2+* recurrent cluster cells by scRNA-seq (>0.75% versus ≤0.75% of tumor cells) (**Table S18**).

To determine the clinical significance of *PLCG2* expression, we performed IHC in an independent cohort of SCLC tumor specimens (N = 31; **Table S17**; **Figure 5E**). Kaplan-Meier analysis revealed worse overall survival in patients with tumors exhibiting high PLCG2 expression (>15% of tumor cells with high PLCG2 intensity; p < 0.002; **Figure 5D**). An adjusted Cox proportional hazards model not only confirmed decreased overall survival, but also showed that high *PLCG2* positivity was a stronger predictor of worse outcome than treatment history, presence of metastatic disease, or SCLC subtype (**Figure S3E**).

Since PLCG2 expression is only proxy to the recurrent cluster 22 cell phenotype, we also assessed within our scRNA-seq cohort if the fractional representation of this subpopulation had prognostic significance. Patients with a subclonal population representing >0.75% of total tumor cells had significantly decreased overall survival relative to others (p = 0.008; **Figure 5E; Table S18**). An adjusted Cox proportional hazards model confirmed worse overall survival with a greater hazard ratio than PLCG2 positivity in the IHC analysis (19.98 vs 8.22), suggesting that while PLCG2 positivity is a strong predictor, it may have reduced specificity for this recurrent tumor population relative to the full transcriptional phenotype (**Figure S3F**). Taken together, these data support that a small stem-like, pro-metastatic subclone with selectively high expression of PLCG2 has a remarkably large prognostic impact across SCLC subtypes.

### Treatment and subtype drive phenotypic shifts in the SCLC immune TME

SCLC is recognized as a particularly immune-cold cancer^46^, and the addition of immune checkpoint blockade to standard-of-care chemotherapy only modestly improves median survival^4,47^. Understanding the specific effects of therapies and the role of both canonical and novel SCLC subtypes in shaping the immune environment will be key to developing effective interventions. However, a comprehensive characterization of the SCLC immune compartment has not been feasible due to limited biospecimen availability and the inaccuracy of low-abundance cell type deconvolution from bulk RNA-seq data.

To address the impact of therapies and SCLC subtype on the immune TME, we analyzed flow cytometry data from our single-cell cohort as well as an independent SCLC cohort (n = 11, **Table S19**). Beyond confirming that the CD45+ population is lower in SCLC than LUAD, our analysis reveals strong reductions specifically in SCLC-N and NEUROD1-positive tumors (**Figures S4A and S4B**). Moreover, we find that the immune population decreases significantly in chemo-treated tumors but is evidently rescued by the addition of immunotherapy (**Figure S4C**).

To assess impacts on cell phenotypes, we pooled immune cells across 21 SCLC samples in our cohort (n = 16,475 cells), using immune cells from LUAD (n = 45,535 cells) and normal adjacent lung (n = 10,934 cells) as a reference (**Figure S4D**), and split the myeloid and T-cell compartments to facilitate cell type annotation (**Figures 6A, S4E–G and S5A–C; STAR Methods**). Our cohort includes 1 SCLC-P, 14 SCLC-A and 6 SCLC-N tumors that are balanced with respect to treatment history (7 untreated, 6 treated with chemotherapy and 8 treated with chemotherapy and immunotherapy). Consistent with flow cytometry (**Figure S4C**), we found that chemotherapy-treated tumors demonstrate lower total immune cell abundance, and that immunotherapy appears to successfully recruit immune cells, particularly T cells, to the tumor (**Figure 6B**). We also see decreased representation of SCLC-N in immune cell phenotypes compared to SCLC-A, suggesting a relatively immune-cold TME (**Figure 6C**). To quantify the impact of treatment and subtype on immune phenotype, we used a measure of entropy based on the k-nearest neighbors of each cell (**Figure 6D**). A comparison of normalized entropies by cell type (**Figures 6D and 6E**) indicates that treatment produces significantly lower entropies (ie, greater phenotypic impact) in the T-cell compartment (CD4+ Tconv, CD4+ Tregs, CD8+ Tmem, CD8+ Texh) compared with subtype effects, consistent with previous reports that treatment heavily influences T-cell phenotypes in other cancers^48–50^, whereas subtype generates significantly lower entropies than treatment for NK cells, cDCs, pDCs, and mast cells (**Figure 6E**). Our results reveal subtype-and therapy-specific differences within the suppressed immune microenvironment of SCLC.

**Figure 6:**
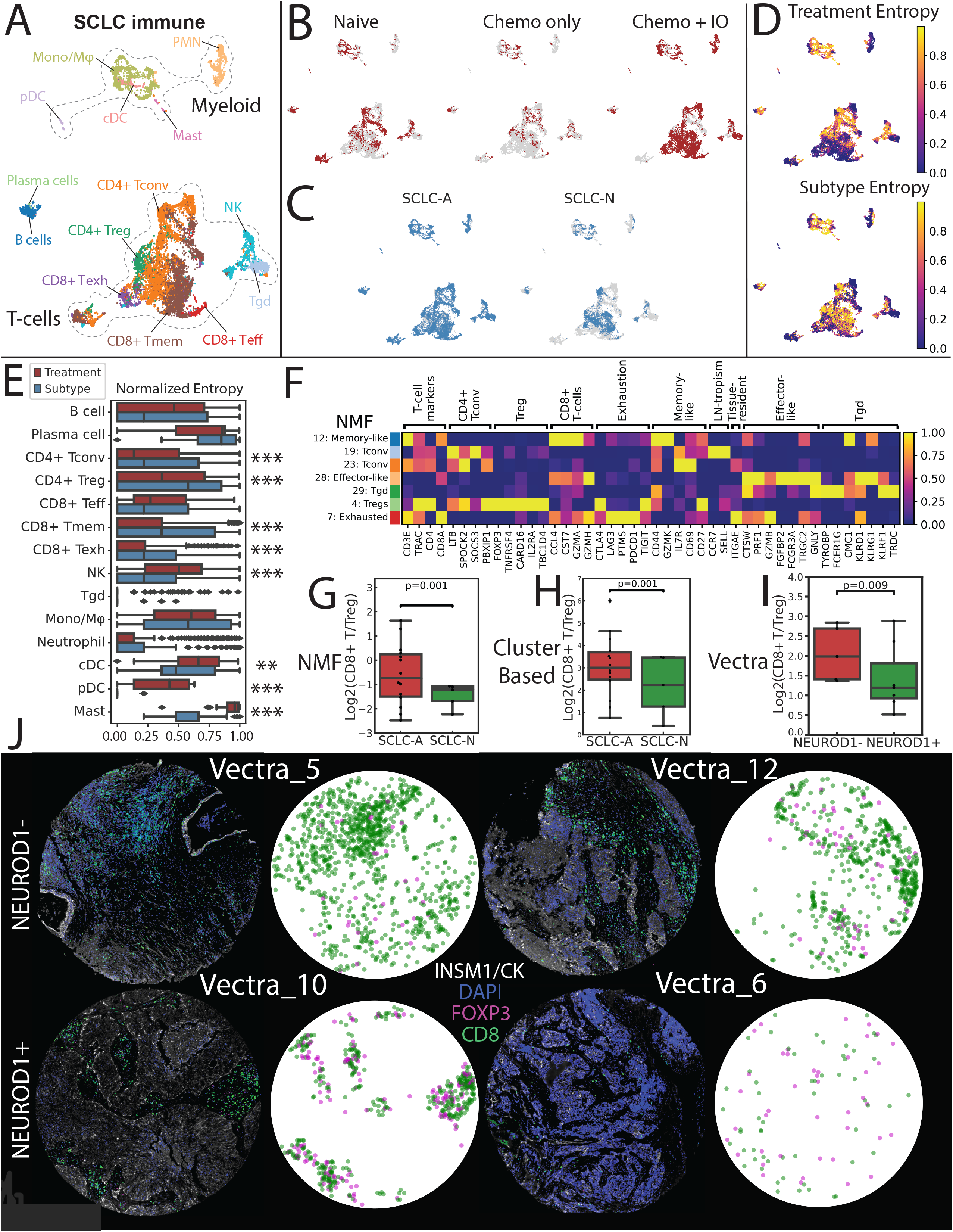
Analysis of therapy and subtype-specific changes in immune phenotype indicate suppressed T-cell activity in SCLC-N. Related to Supplementary Figures S4-5 and Supplementary Tables S4-5. (A) UMAP projections of SCLC immune subsets (**see Figures S4E-G**). Tconv = conventional T-cell; Treg = regulatory T-cell; Teff = effector T-cell; Tmem = memory T-cell; Tgd = γδ T-cell; Mono/Mφ = monocyte/macrophage; PMN = neutrophil; cDC = conventional dendritic cell; pDC = plasmacytoid dendritic cell. (B) UMAP projections of immune cells from SCLC tumors that were treatment-naive or previously treated with either chemotherapy alone or combined with immunotherapy (IO). (C) UMAP projections of SCLC immune cells from tumors that were predominantly SCLC-A or SCLC-N. (D) UMAP projection of entropy based on treatment (top) or SCLC subtype (bottom). Entropy was normalized based on maximum entropy possible given treatment or subtype labels (**STAR Methods**). (E) Barplot showing normalized entropy based on treatment or SCLC subtype per immune subset (**STAR Methods**). *<0.05, **<0.01, ***<0.001. (F) Heatmap of NMF gene loadings for factors associated with T-cell phenotype. Genes are grouped by T-cell function. Each factor is z-scored across genes, and loadings are subsequently scaled from 0 to 1 across factors (**STAR Methods**). Barplot comparing CD8+ Teff/Treg log ratio for based on (G) NMF factors and (H) clusters associated with T-cell phenotype in SCLC-A versus SCLC-N in our single-cell cohort (n=19), adjusted for treatment and tissue site (weighted t-test, **STAR Methods**). (I) Bar plot comparing CD8+ T/Treg log ratio in NEUROD1-versus NEUROD1+ SCLC in an independent cohort with Vectra multiplexed fluorescent imaging (n=12) (weighted t-test, **STAR Methods**). (J) Selected Vectra imaging of NEUROD1-versus NEUROD1+ SCLC (2 samples each). Fluorescent markers include CD8 (cytotoxic T-cells), Foxp3 (Tregs), INSM1/CK7 (epithelial and tumor cells), and DAPI (DNA). CD8 (green) or Foxp3 (pink) positivity of segmented cells are shown (**STAR Methods**).

### Increased T-cell dysfunction in SCLC in metastases and SCLC-N subtype

Next, we assessed how treatment and subtype impact T-cell phenotype in SCLC. Given challenges of low library size and low CD4 transcripts, we included T-cells from LUAD and normal lung to provide increased power for T-cell phenotype annotation. Additionally, to score T-cell phenotypes, we used non-negative matrix factorization^51–53^, which excels in settings of continuous phenotype, where cluster boundaries are less certain (**STAR Methods**). This approach identified 30 factors that associate scores, or loadings, to each cell as well as each gene. For each factor, we associate genes with high loadings to the corresponding cells with high loadings, thus facilitating phenotypic annotation for each cell. Of 30 factors, 7 corresponded to specific T-cell phenotypes: CD4+ regulatory (Tregs, factor 4), CD4+ conventional (Tconv, factors 19 and 23), CD8+ exhausted (Texh, factor 7), CD8+ memory (Tmem, factor 12), CD8+ effector (Teff, factor 28), and CD8+ gamma delta T-cells (Tgd, factor 29) (**Figures 6F and S5A; STAR Methods**). A parallel cluster-based approach to phenotyping confirmed the annotation of these factors (**Figures S5B–E; STAR Methods**).

To assess changes in T-cell phenotype between SCLC subtypes, we compared factor loadings between SCLC-A vs SCLC-N while adjusting for treatment and tissue. We found that compared to SCLC-A, SCLC-N was significantly increased in Treg factor 4 and CD8+ exhausted factor 7, as well as significantly decreased in CD8+ effector-like factor 28 and Tgd factor 29 (**Figure S5F**). A decreased ratio of CD8+ effector to Treg cells has been correlated with poor prognosis in cancer patients in a variety of contexts^54–56^. The ratio of CD8+ effector to Treg factor loadings was significantly decreased in SCLC-N compared to SCLC-A (p = 0.001; **Figure 6G; STAR Methods**). This measure of immunosuppression was consistent with a parallel cluster-based CD8+ effector/Treg ratio (p = 0.001; **Figure 6H**; **STAR Methods**) and validated by Vectra imaging in an independent SCLC cohort (n = 35) with a similarly reduced ratio in NEUROD1+ samples (p = 0.009; **Figures 6I and 6J**; **Table S20; STAR Methods**). Our findings thus identify multiple modes of T-cell dysfunction in SCLC, including the depletion of cytotoxic T-cells in metastases and SCLC-N, and the increase of Tregs in SCLC-N.

### Populations resembling fibrosis-associated macrophages are enriched in SCLC metastases

We next examined the myeloid compartment (n = 2,951 cells), which comprises 13 clusters, including 7 monocyte/macrophage (Mono/Mφ), 4 neutrophil, and 2 dendritic cell populations (**Figures 7A, S6A and S6B; STAR Methods**). Clusters 1, 7, 9, and 12 represent a subset of *THBS1+ VCAN+* Mono/Mφ cells that overexpress genes related to the extracellular matrix (ECM), including *VCAN, FCN1, S100A4, S100A6, S100A8* and *S100A9* (**Figures S6A–D; Table S21, STAR Methods**). This phenotype resembles monocytic myeloid-derived suppressor cells (MDSCs) in mice^57^ and MDSC-like Mφ expressing *THBS1+* S100 proteins in human hepatocellular carcinoma^58^.

**Figure 7:**
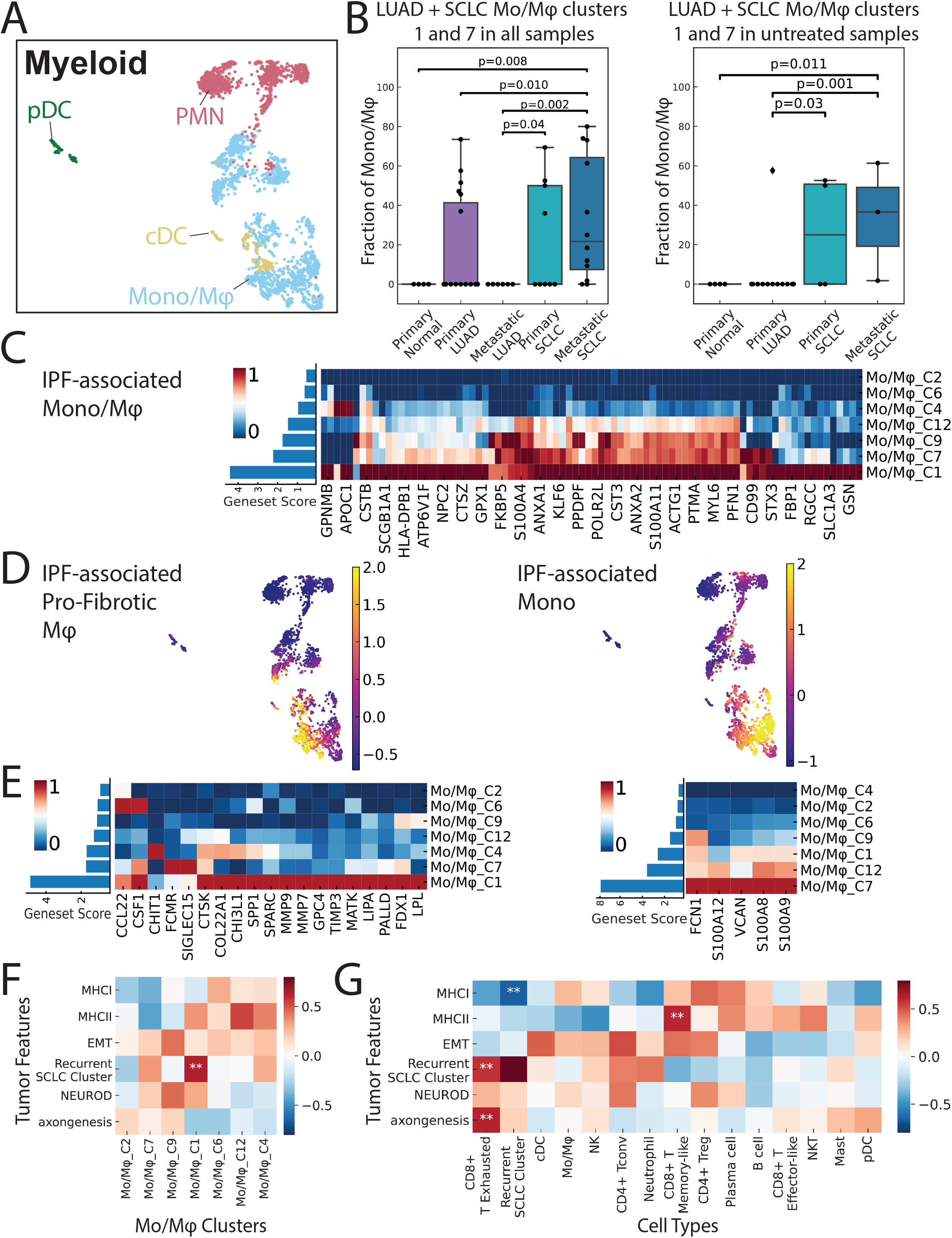
A mixed pro-fibrotic, immunosuppressive Mono/Mφ subset is associated with the recurrent PLCG2-high tumor subclone. Related to Supplementary Figures S6-7 and Supplementary Table S5. (A) UMAP projection of SCLC myeloid cells (n = 2,951 cells) annotated by Phenograph clusters. (B) Box plot showing the proportion of pro-fibrotic Mono/Mφ of the combined myeloid compartment (cluster 6) in different histologies for all samples (n=50) and treatment-naive samples (n=23). (Mann-Whitney test). (C) Heatmap showing the normalized mean expression of markers in the IPF-associated profibrotic macrophage gene signature (n = 143 genes with logFC > 0.3) per Mono/Mφ subsets, and values are subsequently scaled from 0 to 1 across clusters. Left barplot shows the average of z-scored gene expression across the gene signature per cluster. Expression values are imputed using MAGIC (k=30, t=3). (D) UMAP projections of SCLC myeloid cells depicting gene signature scores for IPF-associated pro-fibrotic macrophages and monocytes. (E) Heatmaps showing normalized mean imputed expression of IPF-associated pro-fibrotic macrophage and monocytic gene signatures per SCLC Mono/Mφ cluster. Expression is imputed using MAGIC (k=30, t=3) and scaled from 0 to 1 across clusters. Rows are ordered by average expression across genes per cluster, as depicted in left bar plots. Heatmaps showing covariate-adjusted Spearman’s correlation of SCLC tumor phenotypes with (F) coarse immune cell types and (G) Mono/Mφ subsets. Asterisks denote significant correlation (p < 0.01).

We compared these SCLC-associated Mono/Mφ to those in LUAD. Unsupervised clustering of the full myeloid compartment of SCLC, LUAD, and normal lung reveals that SCLC clusters 1 and 7 group with other LUAD-associated Mono/Mφ cells, suggesting a shared myeloid phenotype in the TME of these cancers (**Figures S7A-C**). Interestingly, this shared phenotype was absent in normal lung and metastatic LUAD and signficantly enriched in SCLC metastases compared to either primary SCLC or primary LUAD. Only treated primary LUAD harbored this myeloid population at all (**Figure 7B**).

We hypothesized that the underlying gene programs of clusters 1 and 7 may facilitate SCLC metastasis. Given that they belong to a Mono/Mφ subset secreting ECM-related proteins, we compared these populations to those in idiopathic pulmonary fibrosis (IPF)^59^. We found that this subset and in particular clusters 1 and 7 closely resemble previously defined IPF-associated macrophage populations (**Figure 7C**). Cluster 1 scores highest for a pro-fibrotic macrophage signature within IPF, and cluster 7 scores highest for a monocytic signature within IPF. (**Figures 7D and 7E**).

Differential expression and pathway analysis (**Figures S7D and S7E, Table S22**) identifies cluster 1 as a *CD14*+ *ITGAX*+ *CSF1R*+ subpopulation that secretes specific pro-fibrotic, pro-metastatic growth factors involved in ECM deposition and remodeling^60^, including fibronectin 1 (*FN1*)^61,62^, cathepsins (*CTSB* and *CTSD*)^63,64^, and osteopontin (*SPP1*)^65,66^, suggesting a role in promoting metastasis.

In addition, cluster 1 overexpressed genes related to immune inhibition, including (1) *SPP1*, implicated in T-cell suppression and tumor immune evasion in colon cancer^67^ and NSCLC^68^; (2) *CD74*, implicated in both immune suppression in metastatic melanoma^69^ and migration inhibitory factor-induced pulmonary inflammation^70^; and (3) *VSIG4*, implicated in macrophage suppression^71^. Collectively, these findings suggest that cluster 1 is a pro-fibrotic, immunosuppressive Mono/Mφ subpopulation that may facilitate SCLC metastasis.

### The recurrent PLCG2-high tumor subclone is associated with a pro-fibrotic, immunosuppressive Mono/Mφ subpopulation and CD8+ T-cell exhaustion

We next hypothesized that the pro-fibrotic, immunosuppressive Mono/Mφ subset may interact with specific tumor subclones to facilitate progression. Although we did not detect an association with canonical SCLC subtypes, we found that cluster 1 is significantly enriched in samples also enriched for the recurrent *PLCG2*-high subpopulation (p = 0.018) (**Figures S7F and S7G; STAR Methods**).

Beyond the association between these myeloid and tumor subsets, we also assessed whether the recurrent *PLCG2*-high tumor subclone correlated with other subpopulations of the TME. We confirmed that the *PLCG2*-high subpopulation is significantly correlated not only with Mono/Mφ cluster 1 (p = 0.004, **Figures 7F**), but also with exhausted CD8+ T cells (p = 0.001, **Figures 7G; STAR Methods**). Notably, the recurrent tumor subclone is also significantly anti-correlated to MHC class I expression of the bulk tumor, suggesting a possible escape mechanism from CD8+ T-cells (**Figure 7G**). Our findings demonstrate that the novel, recurrent SCLC subpopulation exists in an immunosuppressed TME characterized by a pro-fibrotic, immunosuppressive Mono/Mφ and exhausted CD8+ T-cells in the setting of MHC class I downregulation of the bulk tumor.

## DISCUSSION

The paucity of substantial fresh SCLC clinical specimens has been a barrier to capturing SCLC tumor diversity. By optimizing techniques to generate scRNAseq data from both surgical resections and biopsies, we constructed the first extensive single-cell atlas of human SCLC by transcriptomic profiling. With this dataset, we have dissected the biology of SCLC tumor and TME compartments, distinguished features separating LUAD and SCLC subtypes, discovered a novel recurrent SCLC subpopulation with stem-like characteristics, and described its unique TME.

SCLC has been classically considered a homogeneous disease based on a highly consistent microscopic appearance, but recent comprehensive analyses of these tumors has revealed the existence of distinct transcriptomic subtypes^5^ with potential prognostic and therapeutic implications^17,72^. Here, we show that SCLC tumors--particularly SCLC-N--exhibit tumor heterogeneity that exceeds LUAD, highlighting a biological complexity not adequately described by canonical subtyping.

In particular, we found that variant (non-SCLC-A) SCLC subtypes are enriched in local/distant SCLC metastases in humans, with SCLC-N upregulating pathways in EMT, axonogenesis, and metastasis^20^. These findings are consistent with 1) a previously reported mouse model with invasive and metastatic SCLC tumors overexpressing *NEUROD1*^17^ and 2) cell lines with an axonogenic phenotype that correlates with *NEUROD1* expression and promotes metastasis^33^. In contrast to SCLC-A, we found an enrichment of ligand-receptor interactions in SCLC-N. One biphenotypic tumor identified ligands secreted from the SCLC-A compartment that activate Notch, Wnt, TGFβ/Activin and pro-axonogenic signaling in SCLC-N. While this finding in just one tumor should be interpreted with caution, these results reflect preclinical SCLC models describing two discrete intratumoral subpopulations that interact to facilitate metastasis: a high-neuroendocrine subset expressing Notch ligands, and a low-neuroendocrine subset expressing Notch receptors and downstream Notch activation^35,36^. Our data supports this model. Several of these cases support laboratory data suggesting plasticity of SCLC tumors to interconvert between subtypes, and in particular between SCLC-A and -N.

In the search for biological processes that may be universal in SCLC, we found a novel subpopulation that was shared among tumors across subtypes, treatments, and tissue locations. This population demonstrated a pro-metastatic, stem-like phenotype marked by profound *PLCG2* overexpression. Signaling by another phospholipase C gamma family member, *PLCG1*, has been implicated in promoting metastasis in other tumor types^39,40^, suggesting that *PLCG2* overexpression may have biological relevance in SCLC. Compared to primary tumor, we observed increased *PLCG2* expression in local and distant metastases. Direct genetic manipulation validated that *PLCG2* expression correlates with key markers of metastatic potential. Both prevalence of the recurrent tumor subpopulation and *PLCG2* positivity by IHC were stronger predictors of poor survival than treatment, metastasis, or SCLC subtype. These results indicate a novel subtype-independent mechanism that promotes metastasis and worse survival in SCLC, driven by a *PLCG2*-overexpressing subpopulation that exhibits stem-like features.

Analysis of the TME confirmed an immune-cold phenotype in SCLC compared to LUAD with reduced immune infiltrate, consistent with previous reports^73,74^. Within SCLC, we noted increased Tregs and decreased CD8+ T-cells in SCLC-N, indicating greater suppression of adaptive immunity^75–77^ compared to SCLC-A.

Analysis of the myeloid milieu of SCLC revealed a group of immunosuppressive Mono/Mφ resembling IPF-associated macrophages enriched in metastatic SCLC, suggesting a possible tumor-extrinsic mechanism of metastasis through ECM deposition and remodeling. One specific cluster displayed a mixed pro-fibrotic, immunosuppressive phenotype. Interestingly, our analysis of the SCLC cohort identified a constellation of immune and tumor phenotypes (this pro-fibrotic Mono/Mφ, exhausted CD8+ T-cells, and MHC class I depletion of the major tumor subclone), all having significant correlation to the *PLCG2*-high recurrent tumor subclone. These associations raise the possibility that CD8+ T-cells in the TME of the *PLCG2+* tumor subclone are rendered ineffectual due to immunosuppressive Mono/Mφ’s and a lack of MHC class I in the major tumor subclone. This same Mono/Mφ cluster may also provide the fibrotic substrate that facilitates mobility of the pro-metastatic *PLCG2-*high tumor subclone. Targeting these different mechanisms of immune suppression--in SCLC relative to other tumors, among major SCLC subtypes, and within the TME of the recurrent SCLC subclone--may represent a platform for the design of novel subtype-specific and subtype-agnostic immunotherapies.

In conclusion, this atlas of SCLC illustrates how canonical subtypes and a novel *PLCG2*-high recurrent tumor subclone enlist diverse gene programs to create tumor heterogeneity and facilitate metastasis in a profoundly immunosuppressed TME. Our dataset provides further insight into tumor and immune biology in SCLC at single-cell resolution, with potential implications for the design of novel targeted therapies and immunotherapeutic approaches.

## ACKNOWLEDGEMENTS

This work was carried out as part of the NCI Human Tumor Atlas Network (humantumoratlas.org) and supported by NCI U2C CA233284 (DP, CMR), NCI U54 CA209975 (DP), NCI R01 CA197936 and U24 CA213274 (CMR), the SU2C/VAI Epigenetics Dream Team (CMR), the Alan and Sandra Gerry Metastasis and Tumor Ecosystems Center (DP, JMC, OC, IM, LM), the Druckenmiller Center for Lung Cancer Research (CMR, DRJ, MB, TS, AQV), Parker Institute for Cancer Immunotherapy grant (TS); International Association for the Study of Lung Cancer grant (TS), NIH K08 CA248723 (AC), NIH K08 CA245206 (MB), NCI R01 CA217169 and R01 CA240472 (DRJ). We gratefully acknowledge the use of the Integrated Genomics Operation Core, funded by the NCI Cancer Center Support Grant P30 CA08748, Cycle for Survival, and the Marie-Josée and Henry R. Kravis Center for Molecular Oncology. We also acknowledge Kathleen Daniels, David Humphries, Joana Da Silva Leite, Fang Fang, Barbara Oliveira, Magdalena Parys, Mark Kweens and Rui Gardner from the MSKCC Flow Cytometry Core for their invaluable help.

## CONTACT FOR REAGENT AND RESOURCE SHARING

Further information and requests for resources should be directed to and will be fulfilled by the Lead Contact, Charles Rudin.

## STAR METHODS

### Patient cohorts

Patients with NSCLC or SCLC undergoing a surgical resection or tissue biopsy at Memorial Sloan Kettering Cancer Center (MSKCC) were identified and collected prospectively from 2017 to 2019. The current work leveraged four patient cohorts. Single cell RNAseq were performed on 19 clinical specimens with SCLC, 24 clinical specimens with lung adenocarcinoma, and 4 tumor-adjacent normal lung tissue samples (**Table S1**). IHC for subtyping TFs was performed on the SCLC samples as previously described^16^ and reviewed by a pathologist at MSKCC.

IHC and Vectra analyses were performed on a TMA constructed with an additional independent SCLC cohort. 23 cases were amenable for IHC evaluation (**Table S17**) and 27 for Vectra analyses (**Table S20**). For TMA construction, archival formalin-fixed, paraffin-embedded (FFPE) samples were identified and collected retrospectively from SCLC and NSCLC cases between 2007 and 2017. Human kidney samples were used as a positive control in both TMAs.

Flow cytometry analysis of CD45 positive cells was performed on an independent cohort of SCLC patients (**Table S19**) collected prospectively from 2017 to 2019.

Informed consent was obtained for patients in all cohorts (IRB protocol 14-091), reviewed, and accepted by the IRB. Clinical, demographic, pathologic, and molecular data using MSK-IMPACT were identified by retrospective review of the electronic medical record.

### Sample handling

Clinical samples were received in the lab immediately after extraction (Median delivery time±SEM, 0.75±0.72 hours) and processed rapidly (Median ±SEM processing time from delivery until 10x protocol started, 1.75±0.27 hours) to ensure high sample viability and quality for single cell RNAseq.

#### Sample processing: Resection and small biopsies dissociation

Upon delivery to the lab, samples were mechanically/enzymatically dissociated using the tumor dissociation kit (#130-095-929, Miltenyi) and the GentleMACS Octo Dissociator with Heaters (Miltenyi, # 130-096-427). Resection samples were chopped and added to 7.5 mL of enzyme mix in the GentleMACS tube, while core needle biopsies/fine needle aspiration samples were added to 2.5 mL of enzyme mix in the GentleMACS tube. After 15-30 minutes dissociation, depending on sample size and consistency, bigger size samples were filtered with MACS SmartStrainers (70 μm) (Miltenyi, #130-098-462) into 50 mL tubes, and smaller samples were filtered with 35 uM stainer cap FACS tube (Corning # 352235). Then, samples were centrifuged (800g, 1 minute) and supernatant was discarded. Pelleted cells were then stained as indicated below.

#### Sample processing: Pleural effusions cell collection

Upon delivery to the lab, samples were centrifuged at 800g, 10 minutes. The supernatant was discarded and the pellet resuspended in 40 mL of 1X PBS containing 2.5% FBS. Next, 15 mL of Ficoll Paque (GE healthcare, #17-1440-03) was added per tube to two SepMate tubes (STEMCELL Technologies, #85450). Then, 20 mL of pleural fluid was added onto each SepMate tube, slowly, drop by drop, to avoid mixing of the sample and Ficoll, followed by centrifugation at 1200g for 20 minutes at RT. After centrifugation, 15 mL of the upper fluid layer were discarded, and the remaining 5 mL above the dividing plastic surface in the tube were collected, resuspending the cells located in it. Finally, cells were pelleted by centrifugation at 800g, 2 minutes and stained with anti-CD45 antibody and calcein dye as indicated below.

#### Sample processing: staining for sorting and CD45+ composition analyses

Cell pellet was resuspended in 200-3000 uL of Red Blood Cell Lysis Solution (ACK lysis buffer), depending on the pellet size. After incubation for 2 minutes at room temperature the ACK buffer was diluted 10-times with 1X PBS containing 2.5% FBS and pelleted again. Cell pellet was resuspended in 100 uL of 1X PBS + 2.5% FBS, mixed with 5 uL of Human TruStain FcX (Biolegend #422302), 3 uL of PE CD45 antibody (Biolegend # 368510 and 0.1 uL of calcein (1μg/μL, Calcein (Biolegend #425201)), and left for 15 minutes on ice. Stained samples were washed twice with 2 ml of 1X PBS + 2.5% FBS, and finally resuspended in the same buffer supplemented with DAPI dye. Using BD FACSAria (BD Biosciences) or Sony MA900 (Sony) flow cytometers, cells were sorted on DAPI-, Calcein+ (FITC+) to select for live cells. In addition, we sorted CD45+ (immune cells) and CD45-(cell population enriched in tumor cells) populations into separate tubes, and mixed them back in an artificial ratio to balance the compartmental representation (1:5-1:10 ratio, depending on cell availability). To define the percentage of immune cells in each sample, we registered the fration of CD45+ and CD45-in the live cell (DAPI-, Calcein+) population.

#### Sample processing: single-cell RNA-seq

FACS sorted cells were subjected to scRNA-Seq protocol using Chromium (10X genomics) instrument and Single Cell 3’ Reagent Kit (v3). Each sample, containing approximately 3000-8000 cells was encapsulated and barcoded following the manual (CG000183 Rev B). The viability of samples varied between 80-93%, as confirmed with 0.2% (w/v) Trypan Blue staining. The final sequencing libraries were double-size purified (0.6–0.8X) with SPRI beads and sequenced on Illumina Nova-Seq platform (R1 – 26 cycles, i7 – 8 cycles, R2 – 70 cycles or higher). Sequencing depth was between ~160-200 million reads per sample (~50.000 reads per cell).

### Cell lines and PLCG2 overexpression/CRISPR knock out

H82 and DMS-114 were purchased from ATCC and regularly tested for Mycoplasma. Both cell lines were cultured in RPMI 1640 supplemented with 10% FBS.

Lentiviral plasmids were used for PLCG2 overexpression (GeneCopoeia, #EX-A8643-Lv201) and for *PLCG2* CRISPR knock out (Sigma-Aldrich, #HSPD0000031727). Lentiviral particles were produced by standard protocols, transfecting HEK293T cells using JetPrime reagent (Polyplus, #114-15) and concentrated viruses using Lenti-X Concentrator (Takara Bio, #631232) and SCLC cells were transduced at high multiplicity of infection in a spin transduction protocol (Centrifugation of cells at 800xg, 30 minutes with 8ug/mL polybrene).

### Immunoblotting

Protein extraction was performed by pelleting cells and resuspending in cold RIPA buffer (ThermoFisher, #89901) supplemented with phosphatase/protease inhibitors (ThermoFisher, #78446) and incubating for 1 hour on ice. Then, protein extracts were were clarified at 14,000 rpm for 10 min in a refrigerated benchtop centrifuge (Eppendorf, #5340 R). Protein lysates were quantified using a micro BCA protein assay kit (Pierce, #23235) and then diluted with extraction buffer, NuPAGE^®^ LDS sample buffer and reducing reagent (Life Technologies) prior to resolving on 4-12% Bis-Tris gradient gels. Gels were wet-transferred to 0.45 μm Immobilon-FL PVDF membrane (Millipore, #IPFL00010). All primary antibodies were incubated overnight with membranes in TBS Odyssey blocking buffer supplemented with 0.1% Tween-20 (LI-COR, #927-50000), while secondary antibodies (donkey anti-rabbit IRDye 800CW (LI-COR, #926-32213) and donkey anti-mouse IRDye 680LT (LI-COR, #926-68022) were incubated at room temperature with agitation for 1 hr in primary blocking buffer supplemented with 0.01% SDS. Membranes were dried at 37°C and protected from light before imaging (LI-COR; Odyssey Sa). Antibodies for PLCG2 (#3872, Cell Signaling Technology), Beta-catenin (#8480, Cell Signaling Technology), pSMAD1/5 (#9576, Cell Signaling Technology), SMAD1 (#6944, Cell Signaling Technology), SMAD5 (#12534, Cell Signaling Technology), N-cadherin (#14215, Cell Signaling Technology), Vimentin (#5741, Cell Signaling Technology), Twist (#46702, Cell Signaling Technology) and actin (#3700, Cell Signaling Technology) were used. Immunohistochemistry was performed as previously described^16^, using antibodies for ASCL1 (#556604, BD), NEUROD1 (#ab205300, Abcam), POU2F3 (Santa Cruz, #6D1) and PLCG2 (#HPA020100, Sigma-Aldrich).

### In vitro metastasis surrogate analyses

Migration and invasion assays were performed using Cultrex BME Cell invasion assay kit (#3455-096-K, R&D Systems), following manufacturer’s instructions. 50.000 cells were seeded per chamber on day 0 on 0% FBS media, with 10% FBS media in the bottom well, and results were collected on day 4 using a luminescent assay (CellTiter-Glo 2.0 assay, #G9242, Promega). Each experiment was replicated a minimum of three times in independent assays, and the experimental condition was normalized to control condition, which was assigned a value of 1. Analysis of invasion/migration capacity was performed by averaging values in the independent replicates and by performing a two-tailed Student’s t-test to assess for statistical significance.

### Pre-processing of scRNA-seq data

Fastq files from patient samples were individually processed using the SEQC pipeline^12^ based on the hg38 human genome reference and default parameters for the 10x single-cell 3’ library. The SEQC pipeline performs read alignment, multi-mapping read resolution, as well as cell barcode and UMI correction to generate a (cells x genes) count matrix. The pipeline further performs the following initial cell filtering steps: true cells are distinguished from empty droplets based on the cumulative distribution of total molecule counts; cells with a high fraction of mitochondrial molecules are filtered (> 20%); and cells with low library complexity are filtered (cells that express very few unique genes). In addition, we performed additional filtering of empty droplets using the CB2 package with parameter “lower” set at 100 to estimate the background distribution of ambient RNA and an FDR threshold of 0.01 for calling real cells^78^. Putative doublets were removed using the DoubletDetection package (DOI 10.5281/zenodo.2658729). Genes that were expressed in more than 10 cells were retained for further analysis. Combining samples in the entire cohort of samples from SCLC, LUAD, and normal adjacent lung yielded a filtered count matrix of 155,098 cells by 23,628 genes, with a median of 5,654 molecules per cell and a median of 3,041 cells per sample. The count matrix was then normalized by library size, scaled by median library size, and log2 transformed with a pseudocount of 0.1 for analysis of the combined dataset. Principal Component Analysis (PCA) was performed with the top 50 principal components (PCs) retained with 42% variance explained.

### Batch correction of the combined dataset

We performed batch correction in the combined dataset of clinical samples including SCLC, LUAD, and normal adjacent lung using fastMNN with cosine distance applied to the log2-transformed of the library-size normalized count matrix with pseudocount of 1, reduced to top 50 PCs. We favored fastMNN due to the ability to perform hierarchical merging among samples first from the same patient, then from the same histology, with samples containing a greater number of cells merged first. To evaluate the effect of batch correction, we used an entropy-based measure that quantifies how much normalized expression mixes across patients^12^. We constructed a k-nearest neighbors graph (k=30) from the normalized dataset using Euclidean distance and computed the fraction of cells *q_T_* derived from each tumor sample *T* in the neighborhood of each cell *j*. We then calculated the Shannon entropy *H_j_* of sample frequencies within each cell’s neighborhood as:

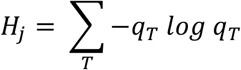

High entropy indicates that the most similar cells come from a well-mixed set of tumors, whereas low entropy indicates that most similar cells derive from the same tumor. This sample entropy was projected on the UMAP (**Figure S1A**). As expected, immune cells generally had the highest entropy consistent with shared phenotypes across tumors, whereas SCLC and LUAD tumor cells had the lowest entropy consistent with increased inter-tumoral diversity. These results indicate that batch effect was corrected while maintaining true biological heterogeneity. Calculating robust clusters

In all cell type compartments, we performed Phenograph clustering^79^ over a range of values for the parameter knn to ensure that subsequent cell typing is consistent. To ensure robustness, we used the adjusted Rand index to evaluate the consistency of clusterings across different k (from 5 up to 100). We then chose knn from the window where the rand index is the consistently highest, indicating stable clusterings. Ultimately we chose knn = 30 for clustering in all cell compartments, with the exception of the T-cell compartment (described below).

### Gene imputation

Given the sparse nature of single-cell sequencing that arises from gene dropout, we used gene imputation using MAGIC (knn = 30, t=3)^80^ when performing knnDREMI calculations (described below) and for visualizing gene expression on both UMAPs and heatmaps (**Figures 2B, S2B, S3B**).

### Single-cell visualization

To visualize single cells of the global atlas as well as epithelial, SCLC, immune, T-cell, and myeloid subsets, we used UMAP projections^81^ to generate lower dimensional representations using knn = 15 and min_dist = 0.3-0.5 (**Figure 1A-C, 2A-B, 4B, 5A-D, 6A, 6D, S1A, S1F, S2A, S3A, S4D-E, S6A, S6C, S7A-C**).

### Coarse cell type identification and subsetting

We used a hierarchical strategy to identify cell types, starting at coarse resolution (epithelial versus immune) and then fine resolution (basal versus NE cell). At the global level, we first performed unsupervised clustering on the batch-corrected count matrix (described below) to identify 58 clusters. Similar to other single-cell studies in lung^82^, we annotated clusters by coarse cell type based on expression of tissue compartment markers (for example, *PTPRC* for immune cells, *EPCAM* for epithelial cells, *COL1A1* for fibroblasts, and *CLDN5* for endothelial cells) (**Figure 1A, Table S23**). We subsetted the data based on these coarse cell types for downstream analysis.

### Measuring inter-patient heterogeneity per cell type

Similar to our evaluation of patient mixing following batch effect, we used an entropy-based measure of inter-patient diversity for each cell type. Here, we use the above PhenoGraph clusters at the batch-corrected global dataset, where each cluster *C* represents a discrete phenotype. We annotate each cluster by cell type (detailed below). To account for any differences in the number of cells per cluster and cell type, we subsampled 100 cells from each cluster 100 times with replacement and calculated the Shannon entropy of patient frequencies *P* in each subsample *H*_*C*_ as:

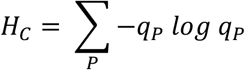

We then compared the distribution of Shannon entropies bootstrapped from clusters between cell types using Bonferroni-adjusted Mann-Whitney U test (**Figure 1D**).

### Cell type annotation of the epithelial compartment

We subsetted the *EPCAM*+ epithelial cells (n=64,301 cells). We projected normalized counts without log transform onto the first 45 PCs selected by detecting the knee-point (minimum radius of curvature in eigenvalues), corresponding to 85.3% variance explained. We identified 38 clusters (described below). We considered a cell cluster to be neuroendocrine based on expression of canonical markers (*CHGA, CHGB, NCAM1*, *SYP*, *ASCL1*, *ASCL2*, *BEX1,* also see **Table S23**). Using this classification, we further divided the epithelial compartment into a neuroendocrine subset (n=54,523 cells) and a non-neuroendocrine subset (n=9,778 cells).

### Cell type annotation of the non-neuroendocrine epithelial compartment

We subsetted the non-neuroendocrine epithelial cells. We projected the normalized counts without log transform onto the first 30 PCs selected by knee-point detection, corresponding to 90.5% variance explained. We then curated multiple recent publications for specific canonical markers for a range of cell types, including epithelial lineages in the lung^82–84^, and liver^85^ (see **Table S23**). Using these cell type-specific gene sets, we first transformed the data by z-score and calculated the average expression of each curated gene set per cell type subtracted from the average expression of a reference set of genes using the score_genes function in scanpy. The subsequent cell type scores were transformed again by z-score, with cell types ultimately annotated by maximum cell type score (**Figure 1A**)

### Tumor cell identification using single-cell SNV and CNV calls

We identify cancer cells in the epithelial compartment by ensuring that all putative tumor cell populations cluster separately from cells derived from normal lung samples. Additionally, we identify tumor cells harboring genomic mutations including single nucleotide variants (SNVs) and copy number variants (CNVs) based on matched bulk DNA-sequencing from MSK IMPACT, downloaded from cBioPortal. To account for the sparsity of scRNA-seq, as well as confounding gene fragments from lysed tumor cells that contaminate normal single-cell droplets, we consider cell clusters to be cancer if they are enriched in reads calling SNVs compared to immune and mesenchymal cells as a negative control, based on Fisher’s p-value adjusted by Bonferroni calculation for multiplicity with a threshold of < 0.05. We reasoned that any cluster with a significant enrichment of variant alleles above a null distribution of normal immune and mesenchymal cells likely represents a cluster of tumor cells.

We also identify CNVs at the single-cell level using InferCNV^86^ using a sliding window of 200 genes, with a diploid mean and standard deviation determined by available normal adjacent tumor samples. We considered any deviations from the diploid mean of at least two standard deviations to be a copy number change.

We noted that the fraction of the genome altered by CNV followed a bimodal distribution across cells, consistent with normal and tumor cells having low and high CNV burden, respectively. We noted that CNV burden was higher in SCLC tumors compared to LUAD (**Figures S1E-F)**, consistent with SCLC having a higher tumor mutation burden^10^. We use two different measures of CNV burden: fraction of the genome changed and Pearson’s correlation between single-cell and bulk CNV profiles, both of which have a bimodal distribution in tumor samples, with a lower peak corresponding to normal stromal cells and a higher peak corresponding to mutated tumor cells. On the other hand, the normal samples have a unimodal distribution that coincides with the normal stromal peak in tumor samples. Based on the bimodal distribution, we identify tumor cells using a threshold of >10% fraction of genome altered and Pearson’s correlation to bulk CNV profile rho >0.2. Of the epithelial cell compartment (n=64,301 cells), clusters that were identified as both tumor and neuroendocrine were therefore subsetted as the SCLC tumor compartment (n=54,523 cells). Epithelial cell clusters identified as tumor but not neuroendocrine (n=7,635 cells) were considered LUAD.

### Differential expression in bulk reference datasets

To facilitate annotation of our single cells by tumor histology and SCLC subtype, we used available reference RNA-sequencing of bulk tumors. These datasets included SCLC subtypes (SCLC-A, SCLC-N, SCLC-P, and SCLC-Y from George, et al.^87^ and Rudin, et al.^88^. We performed differential expression using limma^89^ based on log transcripts per million (TPM) counts (**Tables S2, S3, S4 and S5**). We considered only DEGs with absolute value of log2 fold-change > 1.5 and Benjamini-Hochberg adjusted p-values < 0.05.

### Subtype classification and deconvolution in the SCLC tumor compartment

We aimed to characterize inter-patient tumor heterogeneity of the SCLC tumor compartment within the context of canonical and non-canonical subtypes. To discriminate known SCLC subtypes, we performed feature selection on bulk DEGs between each SCLC-subtype (SCLC-A, SCLC-N, SCLC-P, SCLC-Y) vs rest (described above, **Tables S2, S3, S4 and S5**), and excluded genes from cell cycle, hypoxia, and apoptosis pathways that are non-specific to SCLC subtype and might confound classification. These filtered genes included pathways from REACTOME_CELL_CYCLE_MITOTIC, REACTOME_MITOTIC_G1_G1_S_PHASES, HALLMARK_G2M_CHECKPOINT, HALLMARK_HYPOXIA, HALLMARK_APOPTOSIS downloaded from MSigDB. We projected the normalized counts without log transform onto the first 56 PCs selected by knee-point detection, corresponding to 78.8% variance explained.

We then consider the following semi-supervised classification problem of assigning SCLC subtype. For *N* cells where a subset of *L* cells has known subtype (training data), we must assign the remaining *N-L* cells (test data) the probability of represents subtype *S ∈ {s_1_,s_2_,s_3_} = {SCLC-A, SCLC-N, and SCLC-P}*. We excluded SCLC-Y, as we did not identify any *YAP1*-expressing tumor cells in our SCLC cohort (**Figure 2B**). We want an approach that not only assigns probabilities of each subtype per cell, but is able to deconvolve the phenotype of SCLC tumor cells that reside on a continuum between different SCLC subtypes.

We solve this problem by representing the dataset as an absorbing Markov chain, which is a stochastic model where each event depends only on the state of a previous event. The Markov chain can be represented as a graph where each node is a cell of the set *X = {x*_1_,...,*x*_*N*_}, the set *Y = {y*_1_,...,*y*_*N*_} represents the subtype assignments where *y* is an element in S, and each edge between nodes corresponds to the transcriptional similarity between cells. For cell *x*, we want to calculate the probability of a particular subtype *s*, or *Pr(y*_*x*_ == *s*). Given the training data, we have prior knowledge of the subtype labels of the first *Y*_1:*L*_ = {*y*_1_,...,*y*_*L*_} and must assign subtype labels the the remaining unassigned cells or test data *Y*_*L*+1:*N*_ = {*y*_*L*+1_,...*y*_*N*_}. A simplistic approach to classification of cell *x* would be a “majority vote” of the subset of Y_1:L_ within the knn neighborhood. However, this neighborhood may have skewed or even no available training data due to sampling bias. A better approach would consider distant labeled nodes beyond the immediate neighborhood, connected to *x* through multiple edges in the graph. A “vote” of a labeled node would be weighted by the sum of the edge weights along the path from *x* to the labeled node. However, there may exist multiple paths in the graph from *x* to the labeled node, and so an optimal solution would account for all possible paths at once. An effective way to consider all possible paths is to perform a random walk from *x*, where each step is a traversal from one node to a neighboring node based on relative edge weights. *Pr(y*_*x*_ == *s*) can then be reformulated as the probability that a random walk from *x* will reach a labeled node of subtype *s* first. Rather than simulate this process, Markov absorption probabilities represent an analytical solution for calculating these probabilities, which is implemented in the Phenograph package^79^.

To implement this method, we first must have labeled training data available. To this end, we identify cells that can be confidently assigned to each subtype prior to calculating Markov absorption probabilities. Using reference RNA-sequencing of bulk tumors comparing SCLC subtypes (described above), we used the top 30 overexpressed DEGs per SCLC subtype and calculated the average Z-score over this gene set for each cell. The top 100 highest scoring cells were then used as training examples for Markov absorption calculations.

Next, we constructed a Markov graph from the dataset. We first constructed a diffusion map based on the first 56 PCs to obtain the first 15 diffusion components (DCs) retained by eigengap. Using the Phenograph package, we transformed this diffusion graph additionally into a Jaccard graph between k-neighborhoods, which has been shown to be more robust to noise. The resulting graph represents a Markov chain where we can therefore calculate the Markov absorption probabilities for each unlabeled cell to reach a labeled cell of a given subtype. Based on the resulting probabilities for each subtype, we can then perform a hard classification of SCLC subtype by maximum likelihood, or consider the per-cell probabilities of SCLC-A, SCLC-N, and SCLC-P to be a deconvolution of mixed phenotype that can be readily represented by a 3-coordinate ternary graph, as implemented in the ggtern package^90^(**Figure 2C**).

Of note, hard classification of SCLC subtypes on the UMAP shows that, unlike previously published visualizations of SCLC tumor cells that comprise discrete islands on the layout without clear relation to each other^91^, our feature selection facilitates a visualization that emphasizes patient diversity in the context of SCLC subtypes (**Figure 1A, 2A**).

### Continuity of mixed phenotypes between SCLC-A and SCLC-N

Focusing on SCLC-A and SCLC-N which constituted the majority of our samples, we observed that while most cells were strongly associated with either SCLC-A or SCLC-N, a substantial minority of cells comprised a relatively continuous spectrum of cells from SCLC-A to SCLC-N (**Figure 2C**). This minority (8.9% of cells drawn from 20 samples) comprised a relatively uniform continuum of mixed cell-states with almost any proportion of SCLC-A/N probability. In comparison, cells from our single SCLC-P did not contain any such mixed phenotypes with either SCLC-A or SCLC-N (0.37% of cells). Cells at the transition between SCLC-A vs SCLC-N may represent intermediate subtypes or non-canonical phenotypes.

### Visualizing phenotypic changes along the SCLC-A vs SCLC-N spectrum

For better visualization of SCLC tumor cells along the SCLC-A vs SCLC-N spectrum (**Figure 3B)**, we excluded SCLC-P cells and renormalized the Markov absorption probabilities of SCLC-A and SCLC-N (described above). We ordered the cells by these probabilities from SCLC-A to SCLC-N along the X-axis and colored the corresponding subtype probability on the horizontal color bar. We rescaled marker expression or pathway scores from 0 to 1 along the Y-axis and plotted this value for each cell (grey dots) as subtype probability along the X-axis increasing from SCLC-A to SCLC-N. We calculated pathway scores as the average of Z-scored expression of genes belonging to a pathway. The average trend for each gene marker or pathway was computed by a generalized additive model of 8 splines with spline order 3 using the python package pyGAM (DOI 10.5281/zenodo.1208723).

### Detailed Characterization of Canonical and Non-Canonical Subtypes in the SCLC cohort

To characterize the canonical subtypes in our SCLC cohort, we performed DE analysis between each subtype vs the rest, as well as between the predominant subtypes in our cohort SCLC-A vs SCLC-N, using MAST on the non-imputed count matrix (detailed below, **Tables S7, S8, S9 and S6**). To visualize the gene signatures characterizing each subtype, we plotted the heatmap of imputed gene expression and performed hierarchical clustering to assess the separability and relative similarity of subtype gene signatures (**Figure S2B**). We found typical markers for SCLC-A (*ASCL1, SOX4, STMN2, DOC, STMN2*), SCLC-N (*NEUROD1, ADCYAP1, NRXN1, SSTR2, ID1, ID3, SST, DLK1*), and the one SCLC-P sample (*POU2F3, ASCL2, CD44, MYC, KIT, YBX1*).

Interestingly, sample Ru1108 had a strong subtype probability for SCLC-A but was transcriptionally distinct from the rest of the SCLC-A group (**Figure 2A, S2B**). This sample with wild type *TP53* and *RB1* had high expression of *ASCL1*, *DLL3* and neuroendocrine markers consistent with SCLC-A subtype, but also overexpressed CDK4 and a NSCLC gene signature (average Z-score of the differentially overexpressed genes in NSCLC vs SCLC cell lines from the CCLE database, not shown). Together, our subtype classification demonstrated tumor diversity in canonical SCLC subtypes, but also identified additional non-canonical phenotypes in our cohort, including this *TP53/RB1* wild-type SCLC and the recurrent *PLCG2* tumor subclone (described in main text and below).

### Modeling cell fraction of SCLC subtypes in primary vs metastatic sites

We used several approaches to compare the fraction of tumor cells of different SCLC subtypes in primary lung vs lymph node vs distant metastasis (**Figure S2C**). We performed Dirichlet regression using the DirichletReg R package using *common* parameterization to adjust for treatment status (naive vs chemo-treated vs chemo-immunotherapy treated) and tissue status (primary vs regional lymph node vs distant metastasis). This method tests for differences in cell type composition between groups while accounting for proportions of all other cell subsets. In addition to the multivariate Dirichlet regression, we also used univariate Mann-Whitney as a parallel statistical test to ensure consistency.

### Differential expression of tumor and immune subsets

We performed differential expression for the following comparisons: 1) each SCLC subtype vs rest (**Tables S7, S8 and S9**), 2) SCLC-A vs SCLC-N tumor cells (**Table S6**), 3) Mono/Mφ-A vs Mono/Mφ-B (**Table S21 SAJ**), and 4) each unsupervised cluster vs rest (**Tables S14 and S24**). All differential expression was performed using MAST (version 1.8.2)^92^, which provides a flexible framework for fitting a hierarchical generalized linear model to the expression data. We used a regression model that adjusts not only for cellular detection rate (cngeneson, or number of genes detected per sample), but also tissue status (primary vs LN vs distant metastasis) and treatment status (naive vs chemotherapy vs combined chemo-immunotherapy in the first-line setting):

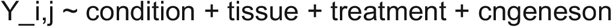

where condition represents the condition of interest and *Yi* is the expression level of gene *i* in cells in cluster j, transformed by natural logarithm with a pseudocount of 1. To homogenize cell sampling per batch, we downsampled such that the cell complexity (i.e. the number of genes per cell) was evenly matched across groups. In particular, we partitioned cells from each cluster into 10 equally-sized bins based on cell complexity and subsampled from each bin to match cell complexity distribution across samples. We downsampled to at most k cells per sample, where k is the median sample size. We verified that the mean expression levels from the full and downsampled datasets were strongly correlated. We considered genes to be significantly differentially expressed for Bonferroni-adjusted p-value < 0.05 and absolute log fold-change > 0.3.

### Filtering ambient RNA from differential expression

To remove candidate DEGs that likely represent ambient RNA, we follow a stepwise, regression-based approach that identifies likely contaminant genes per cell type^93^. For each general cell type (ingroup), expression of each gene is plotted against the expression of that gene in all other cells (outgroup). An initial Loess regression is fitted to the entire dataset. Genes are then binned by expression (number of bins = 25), and the 50 genes with the most negative residuals per bin are then assessed. A second linear regression is fit to genes with negative residuals. Finally, those genes with residuals for the second regression that are < 2 are considered ambient RNA. Likely ambient RNA is colored in red, with known specific markers of other cell types highlighted in red boxes. For instance, *PTPRC* detected in epithelial cells is highly likely to be contaminant RNA from lysed immune cells. We excluded any genes representing ambient RNA from DEGs per cluster or SCLC subtype.

### Identifying gene signatures in single-cell data

Enriched gene pathways were identified using pre-ranked GSEA, as implemented by the R package fGSEA^94^ using 10,000 permutations. Gene ranks were calculated using −log(p-value)*logFC based on MAST^10^ differential expression (described above). To assess enriched pathways in SCLC subtypes and clusters, we used a curated set of pathways from MSigDB v 7.1 (**Table S25**)^95^. To assess enriched pathways in myeloid clusters, we used IPF-related gene sets (see **Table S23**) in addition to HALLMARK and KEGG subset of Canonical Pathways in MSigDB v 7.1^95^. Using the same cutoff as in the original GSEA paper, we considered pathways with Benjamini-Hochberg adjusted p-values < 0.1 to be significant.

### Identifying the recurrent PLCG2+ tumor subclone

A central question beyond canonical SCLC subtypes, was whether there existed any novel tumor phenotypes that are shared across patients. We identified 25 clusters corresponding to distinct SCLC phenotypes. We first assessed whether any of these clusters poorly matched canonical SCLC subtypes and could therefore represent a novel tumor phenotype. Having assigned probabilities for each SCLC subtype *s* for each cell *j* using Markov absorption probabilities *p*_*sj*_ (described above), we identified cells with high uncertainty for any SCLC subtype by calculating the entropy over the cell probabilities for each subtype *U*_*j*_ = Σ_*s*_ *p*_*j*_(s) log *p*_*j*_(s). Cells that have high entropy do not bear clear similarity to any SCLC subtype. We compared the distribution of subtype uncertainties per cluster and found that cluster 22 had significantly higher subtype uncertainties than all other clusters by Mann-Whitney U test, suggesting a novel subtype.

Having identified a possibly novel SCLC phenotype, we next assessed if it arose beyond a single patient. We used a similar approach to assessing inter-patient diversity per cell type (described above), except instead of stratifying by cell type the bootstrapped entropies of patient labels from each cluster, we directly compared the bootstrapped entropies of each cluster versus the rest using Bonferroni-adjusted Mann-Whitney U test. We again identified cluster 22 as the most highly recurrent cluster across patients (**Figure 4A**).

### Recurrent gene markers of the PLCG2+ tumor subclone

To assess the gene program of the recurrent PLCG2+ tumor subclone, we performed differential expression of cluster 22 vs the rest of the tumor cells using MAST (**Table S14**). To assess for recurrence of overexpressed genes across samples harboring cluster 22, we perform differential expression within each tumor sample between cells in cluster 22 and the outgroup. For each gene, we have an adjusted FDR of differential expression in the ingroup vs outgroup, and we calculate a combined p-value **p** by the Edgington’s method to score the recurrence of each gene. In this way, we can avoid pseudoreplication bias that emerges from variably sequenced number of cells per sample^96,97^. We rank the recurrence of each gene by significance −log(**p**) and find *PLCG2* to be the most highly recurrent DEG (**Table S13**).

### Identifying the PLCG2-related gene module

To better characterize the *PLCG2* pathway in the context of SCLC, we used knnDREMI (conditional-Density Resampled Estimate of Mutual Information)^80^ to estimate the functional relationship of *PLCG2* expression to other genes across the dynamic range of expression. To this end, knnDREMI estimates mutual information between two genes by using conditional density instead of joint density. The key feature of knnDREMI is replacing the heat diffusion based kernel-density estimator (KDE)^98^ with a knn-based density estimator^99^, which is robust and scales well in sparse, high-dimensional data. For two genes *x* and *y*, knnDREMI performs a coarse-grained mutual information calculation on a KDE of *p(x,y)*.

First, the KDE is calculated by constructing a knn graph from a fine-grained grid of points. The density at each grid point is computed as:

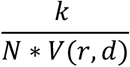

Where *N* is the total number of data points, *k* is the number of nearest neighbors, and *r* is the distance to the *k*th nearest neighbor. *V(r,d)* is then the volume of a d-dimensional ball of radius r:

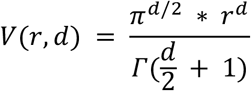

Here, we use d = 2 for considering pairwise relationships between genes and k = 10 to be robust against outliers.

Second, we coarse-grain the KDE to calculate discrete mutual information. While KDE is calculated at fine resolution to smooth and fill in gaps in sparse data, mutual information is calculated over a coarse scale for robustness to noise and any irregularities in partitioning. The conditional density estimate, which is a column-normalized joint density estimate, better captures the functional relationship across the entire dynamic range of expression robust to density sampling.

Finally, we calculate mutual information for gene expression *x* and *y* based on the conditional density estimate. In general, mutual information is defined as

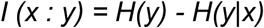

where *H(y)* is Shannon entropy:

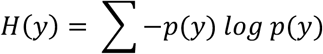

and *H(y|x)* is conditional Shannon entropy:

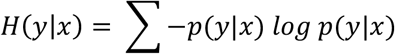

On the other hand, knnDREMI uses the conditional density estimate to calculate mutual information above, which effectively adds another level of conditioning:

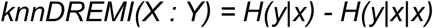

In the SCLC cohort, we identify genes functionally related to *PLCG2* by calculating knnDREMI of each gene *y* conditioned on *x* fixed as *PLCG2* expression. We applied knnDREMI to the MAGIC (t=3) imputed count matrix (described above) and identified genes with the highest knnDREMI > 1. We plotted the z-scored expression of the genes with the highest knnDREMI on a heatmap, ordering column by *PLCG2* expression (top row) (**Figure S3B**). We then performed hierarchical clustering to find three gene modules corresponding to low, intermediate, and high *PLCG2* (**Table S15**).

To identify other pathways associated with the *PLCG2*-high gene module *m*, we calculated for each cell *x* a score *Z*_*m*_, which is the average Z-score of expression for all genes within the *PLCG2*-high gene module. We similarly calculated for each cell a score *Z*_*n*_ the average Z-score of expression for all genes in each pathway *n* from a curated set of MSigDB. We then calculated Pearson’s correlation between *Z*_*m*_ and each *Z*_*n*_ to identify gene pathways that correlate with the *PLCG2*-high gene module. We considered pathways among the top 5% correlated, corresponding to a minimum correlation threshold of ρ = 0.341 (**Figure S3C, Table SAO**). The remaining set therefore represents candidate gene pathways that are also increased in cells that have increased expression of the *PLCG2*-high gene module.

### Survival analysis

To assess the prognostic impact of the recurrent *PLCG2+* subpopulation, we performed survival analyses in our single-cell SCLC cohort and validated these findings in an independent cohort with IHC staining for *PLCG2*. Both cohorts were balanced in number of different covariates, including treatment history and tissue type (**Tables S17 and S18**). For both analyses, we considered samples with extensive-stage ES-SCLC or limited-stage LS-SCLC that recurred (ever had extensive-stage disease). OS was defined as time of biopsy to death or censoring. To separate cohorts under analyses into two subgroups, we used a threshold of (1) at least 0.75% of SCLC tumor cells comprising the recurrent *PLCG2*+ subpopulation as assessed by scRNAseq, or (2) >15% of cells exhibiting high *PLCG2* protein expression (Intensity 3). For our validation cohort with IHC, samples were divided based on *NEUROD1* protein expression into *ASCL1*+ *NEUROD1-* and *ASCL1*(+/−) *NEUROD1*+ subgroups, due to the minimal number of *ASCL1*-*NEUROD1*+ samples and no *ASCL1-NEUROD1-* samples in the cohort.

We then performed Kaplan-Meier (univariate) and Cox proportional hazards (multivariate) survival analysis using the survival R package^100^. In the Cox proportional hazards model, we adjusted for presence of classical vs variant SCLC subtype, treated vs naive, and distant metastasis vs primary/regional lymph node. Our adjusted covariates were dichotomized to ensure a stable fit for the adjusted Cox regression. In general, the corresponding Schoenfeld residuals were invariant to time, but for completeness, we also performed Kaplan-Meier univariate analysis that is independent of the proportional hazards assumption. P-values were calculated using Wald test and were also consistent with bootstrapped p-values.

### Cell-cell interaction analysis

We sought to identify cell-cell interactions among tumor subclones of the same SCLC subtype and between tumor subclones of different subtype. For this analysis, we used CellPhoneDB^101^, which efficiently identifies outlying co-expression of ligand-receptor (L-R) pairs compared to a null distribution generated from permuted cell type labels. We first considered whether tumor-tumor L-R interactions are enriched in SCLC-A vs SCLC-N. Given a list of significant interactions based on CellPhenoDB, we assessed enrichment of interactions using Fisher’s exact test and found that all significant interactions were found in SCLC-N rather than SCLC-A, consistent morphological descriptions of SCLC-N as tightly adherent cells in contrast to SCLC-A (**Figure 3C**).

We then assessed for intratumoral interactions across SCLC-A and SCLC-N subtypes within tumor Ru1215. We identified bidirectional interactions in programs related to neuron development, including axonogenesis and activin signaling^102^. Interestingly, SCLC-A expressed Notch ligands (*JAG2*, *DLK1*^103,104^) and WNT, whereas SCLC-N subclones expressed Notch receptors (*NOTCH1*, *NOTCH2*) and FZD1. These results are supported by preclinical model data showing (1) that combined neuroendocrine and non-neuroendocrine subpopulations within the same SCLC tumor increase metastatic potential^35^; (2) the neuroendocrine subpopulation may promote Notch signaling in the non-neuroendocrine subset^105^; and (3) enrichment of homotypic interactions in neuronal and EMT programs may increase the metastatic potential of the SCLC-N subtype.

### Cell type annotation in the immune compartment

We subsetted the CD45+ immune cells from all SCLC patients (n=16,098 cells). We projected the log2-transformed, normalized counts onto the first 40 PCs based on knee-point detection, corresponding to 26% variance explained. We identified 21 clusters, annotated as B/plasma, T, Myeloid and NK cells using marker genes curated from multiple publications for canonical markers for major immune cell types (including *CD79A, CD3D, CD3E, CD14, ITGAM, ITGAX, MS4A2, SDC1, FCGR3A*; also see **Table S23**). Using these cell type-specific gene sets, we transformed the data by z-score and calculated the average expression of each curated gene set per cell type subtracted from the average expression of a reference set of genes using the score_genes function in scanpy. The subsequent cell type scores were transformed again by z-score and cells annotated by maximum cell type score. Cell type labels were smoothed by cluster after manual inspection to ensure accurate separation of cells.

### Assessing impact of treatment and SCLC subtype on immune phenotype

Similar to our evaluation of inter-tumoral diversity (described above), we use an entropy-based measure for assessing impact of treatment and SCLC subtype on the phenotype of each immune cell type (annotation described below). Here, we consider treatment status (naive vs chemo-treated vs chemo-immunotherapy treated) and SCLC subtype (SCLC-A vs SCLC-N). Of note, immune infiltrate from the single SCLC-P sample was only 50 cells and was disregarded. To account for any differences in the number of cells per cell type, we subsampled 100 cells from each cell type *P* 100 times with replacement and calculated the Shannon entropy of the frequencies for a particular condition of interest *C* (treatment or subtype) within each subsample *H*_*P*_ = *−Σ*_*C*_ *q*_*C*_ *log q*_*C*_. To facilitate comparison between treatment and subtype, we normalized the entropy *H_P_* by the maximum entropy possible, which is −3 log 1/3 for treatment and −2 log 1/2 for subtype. We then compared the distribution of normalized entropies bootstrapped per cell type using Bonferroni-adjusted Mann-Whitney U test (**Figure 6E**).

### Cell type annotation in the T cell compartment

Defining SCLC T-cell subsets was complicated by the relatively lower T-cell infiltrate in SCLC and lower average library size of T-cells in general, both of which can prevent clean separation of subsets based on poorly captured markers like CD4 and CD8. We found that two changes greatly facilitated T-cell phenotyping and separation of CD4+ and CD8+ T-cells. First, we included LUAD and normal lung samples with our SCLC samples to boost the number of T-cells (n=46,140 cells). Second, we z-scored the log2-transformed, normalized counts of each gene, projected onto the first 65 PCs based on knee-point detection, corresponding to 7% variance explained (the relatively lower explained variance is expected given the z-score and log transformation). We then performed annotation of T-cell phenotypes using two following parallel approaches.

#### Non-negative matrix factorization of immune cells

Matrix factorization has been previously used in single-cell analysis^51,106^] and excels in settings of continuous phenotypes which are less amenable for robust partitioning by clustering. In this class of methods, cells and genes are projected into the same lower-dimensional space. The resulting latent factors are associated with weights or loadings for each cell and each gene. These cell and gene loadings can be used to associate gene programs to different cells.

We used non-negative matrix factorization (NMF) implemented in scikit-learn (version 20.0) with default parameters except for tolerance for stopping condition 10^−4^, maximum number of iterations 500, and number of factors *k* = 30. To facilitate comparison across factors, gene loadings were first scaled by standard deviation across genes, then z-scored across factors.

Each factor was then annotated by genes with the highest loadings. By comparing to a reference set of gene markers (**Table S23**), we annotated 7 factors with T-cell phenotypes (2 Tconv, 1 Treg, 1 effector-like, 1 memory-like, 1 exhausted, and 1 Tgd factor). The complete set of NMF loadings are provided in the adata made available for download.

To show robustness of matrix factorization with respect to the number of factors k, we repeated the analysis using *k*=10-50 (data not shown). All 7 T-cell phenotypes could be readily identified with each value of *k*. *k*=30 was ultimately chosen based on knee-point of reconstruction error.

#### Cluster-based approach

In parallel to our factor-based approach, we also performed a cluster-based approach to annotating T-cell phenotypes, similar to our strategy in other cell type compartments. However, given the challenges of T-cell clustering, we performed an additional test of robustness. In addition to confirming robustness of clusterings by adjusted Rand index (described above), we also ensured that clustering was not driven by individual samples. To this end, we repeated clustering with each sample left out and confirmed that the ultimate clustering was robust to dropping samples based on rand index. With these steps, we used Phenograph with knn=40 to ensure robustness, identifying 34 clusters of T-cells pooled from SCLC, LUAD, and normal lung.

We then performed differential expression between each cluster and the rest (described above) and compared DEGs to curated markers of T-cell phenotype (**Table S23**) (**Figure S5B-C**). Finally, we confirmed agreement of our cluster-based cell typing with NMF factors, by calculating the mean cell loadings of each T-cell annotated factor within each cluster (**Figure S5D)**and each cluster-based cell type (**Figure S5E**). Having successfully identified T-cell subsets at the combined level, we confirmed that these annotations restricted to SCLC were also consistent with known gene markers (**Table S23**).

### Comparing T-cell phenotype between SCLC-A vs SCLC-N

To analyze the phenotypic shifts in T cell compartment across SCLC subtypes, we considered NMF factors associated with T-cell phenotype (described above). Using NMF, we compared the distribution of factor loadings across T cells in SCLC-A and SCLC-N. To ensure that factors are assessed on the same scale, we first log2-transformed cell loadings with a pseudocount of 0.0001, shifted the minimum of each factor to 0, and scaled each factor by standard deviation across cells. We accounted for the effect of treatment and tissue site by fitting a linear model between the factor loadings and the treatment and tissue status of cells. We then performed a Bonferroni-adjusted two-sample t-test on the residuals of the factor loadings (**Figure S5F**). We used tissue status (primary vs LN vs distant metastasis) and treatment status (naive vs chemo vs chemo-IO in the first-line setting) as covariates in the model.

### Analysis of CD8+ T cell/Treg ratio in SCLC subtypes

As a measure of immune response in tumor-infiltrating lymphocytes that can be readily calculated from both scRNA-seq and Vectra imaging platforms, we used the ratio of CD8+ T-cells to Tregs in SCLC-A versus SCLC-N. We first compared the ratio of CD8+ T effector/Tregs phenotypes using NMF factors (described above). Specifically, we compared the ratio of the averaged loadings of factor 28 (effector-like) and factor 4 (Tregs) across T cells per sample in SCLC-A and SCLC-N. We accounted for the effect of treatment and tissue site by fitting a linear model between the ratio of CD8+ T effector factor loading/Treg factor loading and the treatment status and tissue site of the samples (similar to correlation analysis described below), and comparing the model residuals. We accounted for the difference in numbers of cells collected per sample using a weighted one-sided t-test (as implemented by ttest_ind in the python library statsmodels). Within each SCLC subtype, the weight of the i-th sample was given by:

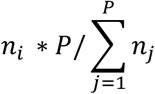

with *n*_*j*_denoting the total number of T cells in patient *i* and *P* being the total number of patients in that group (SCLC-A or SCLC-N). We calculated FDR by generating a null distribution using a permutation test on cell type labels. We also performed Goodman-Kruskal’s test as a parallel statistical test to ensure consistency. To ensure the results are not driven by individual samples, we performed leave-one-sample-out cross-validation and verified that the result remains significant for every case.

We verified the same difference in factor-based ratio of CD8+ T-cell/Treg abundances between SCLC-A vs SCLC-N using several approaches. We first performed the same analysis by using cells labeled with cluster-based T-cell phenotyping (described above).

Finally, we used Vectra imaging to validate these findings. We restricted analysis to 12 treatment-naive, primary SCLC samples. We then compared the ratio of CD8+ T cells/Tregs in NEUROD1-and NEUROD1+ subtypes to quantify the immune response of tumor-infiltrating lymphocytes (described below).

### Analysis of Vectra imaging

#### Batch normalization

To compare different markers across samples, we normalized intensity values of each marker. We first applied a Gaussian kernel with σ=3 to smooth intensity over the target image. We considered the maximum intensity value M of a marker in a given sample to be an initial value for intensity normalization. We then assessed the distribution of maximum intensity values of each marker across samples, which generally follows a bimodal distribution. This bimodal distribution allows for an intensity threshold that readily separates signal from noise. We therefore considered the filtered distribution of intensities greater than this threshold. Finally, we constrained the value for intensity normalization M to be greater than the minimum but less than the maximum of the filtered intensity values across samples.

#### Noise removal

We used the following procedure to remove noise introduced by non-specific staining in our fluorescence multiplexed imaging data. First, we applied a median filter with size 2 to remove outliers, and then a Gaussian kernel with σ=1 was applied to smooth the image. We automated remaining noise removal using either Otsu or Triangle thresholding. For a specific channel, if the 80th percentile intensity is ⋧5, we use the Otsu method. Otherwise we used the Triangle method. To guide automatic noise removal, we manually set a lower boundary (to remove obvious noise) and an upper boundary (to retain obvious signal) per sample. We then combine batch normalization and noise removal to generate a quality check report to further guide preprocessing. This initial automation facilitates manual correction of parameters for image processing.

#### Single-cell instance segmentation

To obtain single-cell information, we adapted Mask R-CNN (https://github.com/dpeerlab/MaskRCNN_cell), a deep learning framework for object instance segmentation to perform cell instance segmentation on our multiplexed imaging data. This model generates bounding boxes and segmentation masks for each instance of an object in the image. We optimized the parameters of this framework for the single-cell segmentation task, characterized by high object density, small but consistent object size. To avoid cropping TMA images into small pieces and cutting cells overlying boundaries into two, we developed seamless stitching features that allow segmentation on very large images. To generate the training data, we manually annotated 24 sample images with nuclear and cell membrane markers (DAPI, CD8, FOXP3, INSM1, et al.) at ImageJ^107^. Training images were augmented by random horizontal flips, random vertical flips, random rotation, random gaussian blur, random zoom in and zoom out, random brightness changes, and random shear. Training was performed using a step per epoch of 1000 and was run for 10 epochs for heads layers thraning and 30 epochs on all layers. To segment images of interests, we visualize the images with the same color pattern that was used in training.

#### Cell typing

The image dataset was subject to segmentation, normalization, and noise-removal, as described above, yielding a 7-dimensional single-cell protein marker expression profile with sum of marker expression, expression area, cell size et al information. Cells with low nuclear area (lower than 16 pixels) were removed prior to analysis. A marker was considered positive when the average expression (total expression divided by cell size) is above 0.1 (0.06 for lowly expressed markers Perforin, FOXP3, CTLA4) and expression area is above 4 pixels. For markers that do not co-express, we classified cells into double-negative, 1 marker positive only and 2 markers positive only, based on the distribution of average expression.

### Cell type annotation in the myeloid compartment

We subsetted the myeloid cells from SCLC patients (n=2,951 cells). We projected the log2-transformed, normalized counts onto the first 50 PCs based on knee-point detection, corresponding to 30% variance explained. We identified 13 clusters, including 7 clusters of monocyte-derived myeloid cells, 4 clusters of granulocyte-derived myeloid cells, and 2 clusters of dendritic cells (**Figure S6A-B**). To annotate myeloid subsets, we identified DEGs between each cluster vs the rest and compared these genes to curated markers of each myeloid subset (**Table S23**).

We sought to further characterize Mono/Mφ in the SCLC cohort. We performed hierarchical clustering of Mono/Mφ clusters using correlation distance based on the mean gene expression profile of each cluster (**Figure S6A**). Based on the resulting dendrogram, we defined an optimal distance threshold that merged clusters into two general groups Mono/Mφ-A and Mono/Mφ-B (**Figure S6B**). Differential expression between each between Mono/Mφ-A vs Mono/Mφ-B was performed using MAST adjusted for treatment and tissue type (**Figure S6C-D**). In line with a more immature phenotype, Mono/Mφ-A overexpressed markers typical of blood monocytes that are also related to the extracellular matrix (ECM), including *VCAN, FCN1,* and S100 proteins (*S100A4, S100A6, S100A8, S100A9*), as well as AP1 transcription factors (*JUNB*, *FOS*, *DUSP1*). *THBS1* and *VCAN* expression in Mono/Mφ-A have also been noted in monocytic myeloid-derived suppressor cells (m-MDSCs) in mice, with *VCAN* driving EMT in lung metastases^57^. On the other hand, Mono/Mφ-B was relatively more heterogeneous (**Figure S6C**) and displayed non-overlapping, specialized gene programs (**Figure S6D**), suggesting a more differentiated group of tumor-associated macrophages (TAMs). Specifically, cluster 4 overexpressed C1 complex (*C1QA, C1QB, C1QC*) and reactive oxygen species (*SOD2, GPX1*) cluster 6 overexpressed inflammatory cytokines (*IL1B*, *CXCL8*, *CCL3*, *CCL4*). The dichotomous Mono/Mφ-A and Mono/Mφ-B phenotypes mirror tumor-enriched macrophages in other cancers, including MDSC-like *THBS1*+ macrophages (expressing S100 proteins) and TAM-like *C1QA*+ macrophages in hepatocellular carcinoma^58^.

### Detailed characterization of pro-fibrotic Mono/Mφ cluster 1

Given the high expression of ECM-related genes in Mono/Mφ-A, we compared our dataset to gene signatures from a single-cell atlas of IPF^59^ and found that cluster 1 stood out as having an outlying pro-fibrotic signature as well as increased inflammatory macrophage signature (**Figures 7B, 7D, 7E**). Differential expression and pathway analysis (**Figures S7C and S7D**) identifies cluster 1 as a *CD14*+ *ITGAX*+ *CSF1R*+ subpopulation. While cluster 1 shared monocytic features with the rest of Mono/Mφ-A, it also overexpressed scavenger receptor (*MARCO*, *MSR1*, *CD36*, *CD68*, *CD163*) and scavenger binding protein (*APOE, APOC1*) genes, suggesting that cluster 1 represents a monocyte-derived but tissue-enriched myeloid subset. In addition, cells from this cluster express secrete pro-fibrotic, pro-metastatic growth factors involved in ECM deposition and remodeling^60^, including *FN1*^61,62^, cathepsins (*CTSB* and *CTSD*)^63,64^, and *SPP1*^65,66^, suggesting a role in promoting metastasis. In addition, cluster 1 overexpressed genes related to immune inhibition, including (1) *SPP1*^67^ and NSCLC^68^]; (2) *CD74*^69,70^; and (3) *VSIG4*^71^. While this cluster did overexpress some markers of immune activation like C1 complex compared to other Mono/Mφ-A clusters (**Figure S7D**), the expression was far decreased compared to activated TAM-like Mono/Mφ-B (**Figure S6E**).

### Correlation analysis of immune subset abundance and tumor phenotypes

We aimed to identify significant correlations between any immune subset and tumor phenotype in SCLC while adjusting for any clinical covariates. To this end, we first consider cell abundance *X* and cell abundance *Y* of interest, as well as clinical covariates *Z*. We fit separate linear regression models between *X* and *Z*, and between *Y* and *Z*. We then compute the Spearman’s rank correlation between model residuals of *X* and *Y*. For this analysis, we adjusted for tissue status (primary vs lymph node vs distant metastasis) and treatment status (naive vs chemo vs chemo-immunotherapy in the first-line setting). We calculated the false discovery rate (FDR) by generating a null distribution using a permutation test on the cell type labels. To test robustness, we performed a leave-one-sample-out validation and confirmed that the result remains significant even after excluding any sample.

## DATA AND SOFTWARE AVAILABILITY

Software and tools used for the enclosed data analysis will be provided open source at http://github.com/dpeerlab. In collaboration with the NIH-funded HTAN Data Coordinating Center (U24), single-cell analysis at time of publication will be made available as an interactive, online platform for independent visualization and analysis.

## SUPPLEMENTARY FIGURE LEGENDS

**Supplementary Figure S1. Related to Figures 1 and 2 and to Supplementary Table S1.**

(A) UMAP projection of the SCLC cohort at the global level (n=155,098 cells) colored by cell type (top) and by Shannon entropy of patients in a k-nearest neighborhood of each cell (k=30, **STAR Methods**) (bottom).

(B) Oncoprint showing mutational information of SCLC samples subjected to MSK-IMPACT bulk targeted DNA sequencing, arranged by genes (rows, sorted by most to least recurrent) and patients (columns). Point mutations are depicted by bars, copy number changes by full squares. Bar plot shows tumor mutation burden (TMB). Annotations at bottom include early-line and later-line treatment history, with treatment order specified in the order of full square, thin rectangle, and small square. Other annotations include smoking status, tissue, stage, disease status, and SCLC subtype of the major subclone.

(C) TMB of SCLC samples, grouped by subtype.

(D) Single-cell CNVs, inferred from scRNA-seq data using InferCNV^86^), arranged by cells (rows) and genomic position (column). Cells are grouped by SCLC, LUAD, and normal lung.

(E) CNV burden of SCLC, LUAD, and normal lung samples. CNV burden is estimated by percent of genome altered by CNV (left) or Pearson’s correlation to CNV profiles of the matched MSK-IMPACT data where available (right). Tumor cells were called based on a threshold of either percent of genome with CNV > 0.1, or bulk correlation ρ > 0.2 (**see STAR methods**).

(F) Clusters comprised by the SCLC (left) or NSCLC (right) samples in our cohort. Each cluster was assigned a different color.

(G) Immunohistochemistry data available for staining of *ASCL1*, *NEUROD1*, *POU2F3* and *YAP1* transcription factors in the SCLC cohort.

(H) Statistics of SCLC subtype representation in samples. Top: SCLC subtype uncertainty per cell as measured by Shannon entropy of SCLC subtype probabilities. Bottom: SCLC subtype fractions in each sample, as determined by maximum likelihood of Markov absorption probabilities.

**Supplementary Figure S2. Related to Figures 2 and 3 and to Supplementary Table S1.**

(A) UMAP of sample Ru1215 showing admixed SCLC-A and SCLC-N subtyping, supported by MAGIC-imputed *ASCL1* and *NEUROD1* expression (**STAR Methods**).

(B) Single-cell gene signatures of SCLC subtypes, with scaled expression (z-score) of differentially expressed genes (DEGs) per SCLC subtype (rows) for each cell (columns). Representative DEGs including canonical TFs are labeled. Top dendrogram produced by hierarchical clustering using Pearson’s correlation as a distance metric with complete linkage. Top annotations include patient and probability of SCLC subtype.

(C) Bar plot of frequency of SCLC subtype per tissue site (primary lung vs regional lymph node (LN) vs distant metastasis). The frequency of each subtype per sample was first determined and then summed by tissue site. A final frequency was normalized by number of samples to produce the stacked bar plot. In this way, each sample was considered with equal weight. SCLC-A was significantly enriched in primary lung whereas non-SCLC-A subtypes are enriched in nodal and distant metastases (Dirichlet regression, p<3.4×10^−8^, **STAR Methods**).

(D) Gene programs significantly enriched in SCLC-A (red bars) or SCLC-N (green bars) using Gene Set Enrichment Analysis (Benjamini-Hochberg adjusted FDR < 0.05, absolute value of normalized enrichment score (NES) > 1) (**Table S6**).

(E) Intratumoral interactions between SCLC-A and SCLC-N subclones in sample Ru1215. Dot size = % cells expressing gene; dot color = mean expression log_2_(X+1), where X is the normalized count. Arrows indicate directionality of interaction (Ligand→Receptor). Bidirectional arrows indicate interactions for which genes can serve as both ligand and receptor.

**Supplementary Figure S3. Related to Figures 4 and 5 and to Supplementary Table S2.**

(A) Unimputed *PLCG2* expression vs log library size in recurrent cluster 22. Library size is transformed by natural log with pseudocount of 1. *PLCG2* expression is mapped on the SCLC UMAP in units of log2(X+1) where X are counts (right).

(B) Scaled expression (z-score, imputed by MAGIC with k = 30, t = 3) of genes with high knnDREMI conditioned on *PLCG2* > 1 (rows), with cells ordered by *PLCG2* expression (columns). For visualization, expression was smoothed over the ordered cells with a rolling window of 100 cells. Hierarchical clustering of genes on the unsmoothed, imputed expression was performed with complete linkage and Pearson correlation as a distance metric, identifying 3 gene modules that predict low, medium, and high *PLCG2* (purple, gray, and yellow respectively) (**Table S15**). Top annotations include *PLCG2* expression and SCLC subtype.

(C) Pathways with average z-scores of gene expression that are highly correlated (>95th percentile, indicated by red line) with the average z-score of gene expression in the high-*PLCG2* gene module (**STAR Methods, Table S16**).

(D) Western blots showing PLCG2 overexpression in H82 and downregulation in DMS114, as well as the activation of Wnt and BMP signaling pathways and expression of indicated markers (PLCG2 = *PLCG2* overexpression, sgPLCG2 = CRISPR knock out).

(E) Multivariate analyses (Cox regression) for overall survival in samples included in an independent cohort of SCLC samples, including the comparison between tumors with >15% cells with high PLCG2 high intensity (defined as intensity >2) versus the rest.

(F) Multivariate analyses (Cox regression) for overall survival in samples included in the single cell cohort, including the comparison between tumors with >0.75% composition of high *PLCG2*-expressing recurrent cluster versus the rest.

**Supplementary Figure S4. Related to figure 6 and to Supplementary Table S1.**

(A) Boxplot of CD45+ infiltrate in LUAD vs SCLC-A vs SCLC-N samples in our single-cell cohort (Mann-Whitney test).

(B) Boxplot of CD45+ infiltrate in ASCL1+ NEUROD1-vs ASCL1+ NEUROD1+ samples in an independent IHC cohort of SCLC tumors (Mann-Whitney test). No ASCL1-NEUROD1+ samples were available.

(C) Boxplot of CD45+ infiltrate in untreated tumors and tumors treated with first-line chemotherapy alone or chemotherapy plus immunotherapy (IO) (Mann-Whitney test).

(D) UMAP projections of the immune subsets of SCLC with LUAD and adjacent normal lung reference at the global level (left, n=73,047 cells), T-cell compartment (middle, n=46,140 cells), and myeloid compartment (right, n=14,072 cells).

(E) UMAP projection of all immune cells from 21 SCLC samples (n = 16,475 cells), annotated by Phenograph clusters.

(F) Dot plots show relative frequency and expressing cells, and mean normalized expression of canonical immune cell type markers per Phenograph cluster.

(G) Dot plots show relative frequency and expressing cells, and mean normalized expression of canonical immune cell type markers per coarse cell type.

**Supplementary Figure S5. Related to figure 6 and to Supplementary Table S5.**

(A) Heatmap of gene loadings of all 30 NMF factors, grouped by T-cell phenotype. The weight of each factor loading is scaled across factors from 0 to 1.

Dot plots show gene markers of each cluster (B) and T-cell phenotype (C). Dot size = % cells expressing gene; dot color = mean expression scaled from 0 to 1.

Heatmap of cell loading of select NMF factors averaged within each cluster (D) and T-cell phenotype based on clusters (E). The resulting averaged loading was z-scored across factors (**STAR Methods**).

(F) Bar plot comparing scaled NMF cell loadings for factors related to T-cell phenotype between SCLC-A and SCLC-N (two-sample t-test adjusted for treatment and tissue site, **STAR Methods**). *** p <0.001.

**Supplementary Figure S6. Related to figure 7.**

(A) UMAP projection of SCLC myeloid compartment, annotated by the different clusters identified.

(B) Heatmap of Pearson’s correlation between the mean expression profile of Mono/Mφ subsets in SCLC, ordered by hierarchical clustering.

(C) UMAP projection of SCLC myeloid compartment, annotated by tumor-associated subsets of *THBS1+ VCAN+* Mono/Mφ-A representing a more immature, monocytic, MDSC-like phenotype and Mono/Mφ-B representing more activated, differentiated phenotypes (**STAR Methods**).

(D) Volcano plots show the top DEGs in Mono/Mφ-A and Mono/Mφ-B (**STAR Methods, Table S21**) highlighting significantly differentially expressed secreted factors (left) and all genes (right).

(E) Dot plot of SCLC myeloid subsets and expression of relevant markers. Myeloid subsets are ordered based on hierarchical clustering of the first 50 principal components (**STAR Methods**). Dot size = % cells expressing gene; dot color = mean expression scaled from 0 to 1.

**Supplementary Figure S7. Related to figure 7.**

UMAP projections of combined myeloid compartment (SCLC, LUAD, and normal adjacent lung), annotated by (A) tumor histology, (B) Mono/Mφ clusters identified from the SCLC-restricted myeloid compartment, and (C) clusters identified from the combined myeloid compartment.

(D) Volcano plot of differentially expressed genes in SCLC Mono/Mφ cluster 1 of the SCLC myeloid compartment versus other Mono/Mφ subsets (**Table S22I**).

(E) Gene programs significantly enriched in SCLC Mono/Mφ cluster 1. Bar plot of NES from GSEA for each pathway. Significantly enriched pathways have FDR < 0.25 and NES > 1 (**Table S24)**.

(F) Boxplots showing the composition of Mono/Mφ clusters stratified by PLCG2+ tumor subclone fraction > 0.75%. We show significant overrepresentation of Mono/Mφ cluster 1 in samples harboring the recurrent PLCG2-high tumor subclone.

(G) Bar plot showing robustness of covariate-adjusted Spearman correlation p-values using leave-one-sample-out cross-validation.

## SUPPLEMENTARY TABLE LEGENDS

**Supplementary Table S1.** Clinical characteristics of samples analyzed by single-cell RNA-seq.

**Supplementary Table S2.** Differentially expressed genes of SCLC-A versus rest in bulk RNA-seq data from George, et al.^87^ and Rudin, et al.^88^ using limma.

**Supplementary Table S3.** Differentially expressed genes of SCLC-N versus rest in bulk RNA-seq data from George, et al.^87^ and Rudin, et al.^88^ using limma.

**Supplementary Table S4.** Differentially expressed genes of SCLC-P versus rest in bulk RNA-seq data from George, et al.^87^ and Rudin, et al.^88^ using limma.

**Supplementary Table S5.** Differentially expressed genes of SCLC-Y versus rest in bulk RNA-seq data from George, et al.^87^ and Rudin, et al.^88^ using limma.

**Supplementary Table S6.** Differentially expressed genes comparing SCLC-A versus SCLC-N in single-cell RNA-seq of the SCLC tumor compartment using MAST.

**Supplementary Table S7.** Differentially expressed genes comparing SCLC-A versus rest in single-cell RNA-seq of the SCLC tumor compartment using MAST.

**Supplementary Table S8.** Differentially expressed genes comparing SCLC-N versus rest in single-cell RNA-seq of the SCLC tumor compartment using MAST.

**Supplementary Table S9.** Differentially expressed genes comparing SCLC-P versus rest in single-cell RNA-seq of the SCLC tumor compartment using MAST.

**Supplementary Table S10.** Pathway enrichment in SCLC-A versus SCLC-N in the SCLC tumor compartment using GSEA.

**Supplementary Table S11.** Pathway enrichment in SCLC-P versus rest in the SCLC tumor compartment using GSEA.

**Supplementary Table S12.** Pathway enrichment in the recurrent, *PLCG2*-high SCLC cluster versus rest in the SCLC tumor compartment using GSEA.

**Supplementary Table S13.** Differentially expressed genes that are recurrently overexpressed in the recurrent tumor subclone across samples, ranked by the Bonferroni-adjusted Edgington’s combined p-value.

**Supplementary Table S14.** Differentially expressed genes of SCLC recurrent subclone (cluster 22) versus rest in single-cell RNA-seq using MAST.

**Supplementary Table S15.** Gene modules with high knnDREMI conditioned on *PLCG2*, divided by low, medium, and high *PLCG2* expression.

**Supplementary Table S16.** Pathways with average z-scores of gene expression correlated with the average z-score of gene expression in the high-*PLCG2* gene module.

**Supplementary Table S17.** Survival data and clinical covariates of an independent tissue microarray stained by immunohistochemistry for PLCG2.

**Supplementary Table S18.** Survival data and clinical covariates of samples analyzed by single-cell RNA-seq and stratified by fraction of the recurrent, *PLCG2*-high SCLC cluster.

**Supplementary Table S19.** Clinical characteristics, CD45+ percentage, and ASCL1/NEUROD1 positivity on immunohistochemistry for an independent SCLC cohort analyzed by flow cytometry.

**Supplementary Table S20.** Clinical characteristics and ASCL1/NEUROD1 positivity on immunohistochemistry for an independent SCLC cohort analyzed by Vectra.

**Supplementary Table S21.** Differentially expressed genes of Mono/Mφ-A vs Mono/Mφ-B in single-cell RNA-seq using MAST.

**Supplementary Table S22**. Differentially expressed genes of Mono/Mφ cluster 1 vs other Mono/Mφ subsets in single-cell RNA-seq using MAST.

**Supplementary Table S23.** GMT file of markers used for cell type annotation, curated from literature.

**Supplementary Table S24.** Pathway enrichment in Mono/Mφ cluster 1 vs other Mono/ Mφ subsets using GSEA.

**Supplementary Table S25.** GMT file of curated set of pathways for GSEA analysis of SCLC subtypes and clusters.

