## Supplementary figures and images for "Single cell profiling reveals novel tumor and myeloid subpopulations in small cell lung cancer"

### Supplemental Figures S1-S7

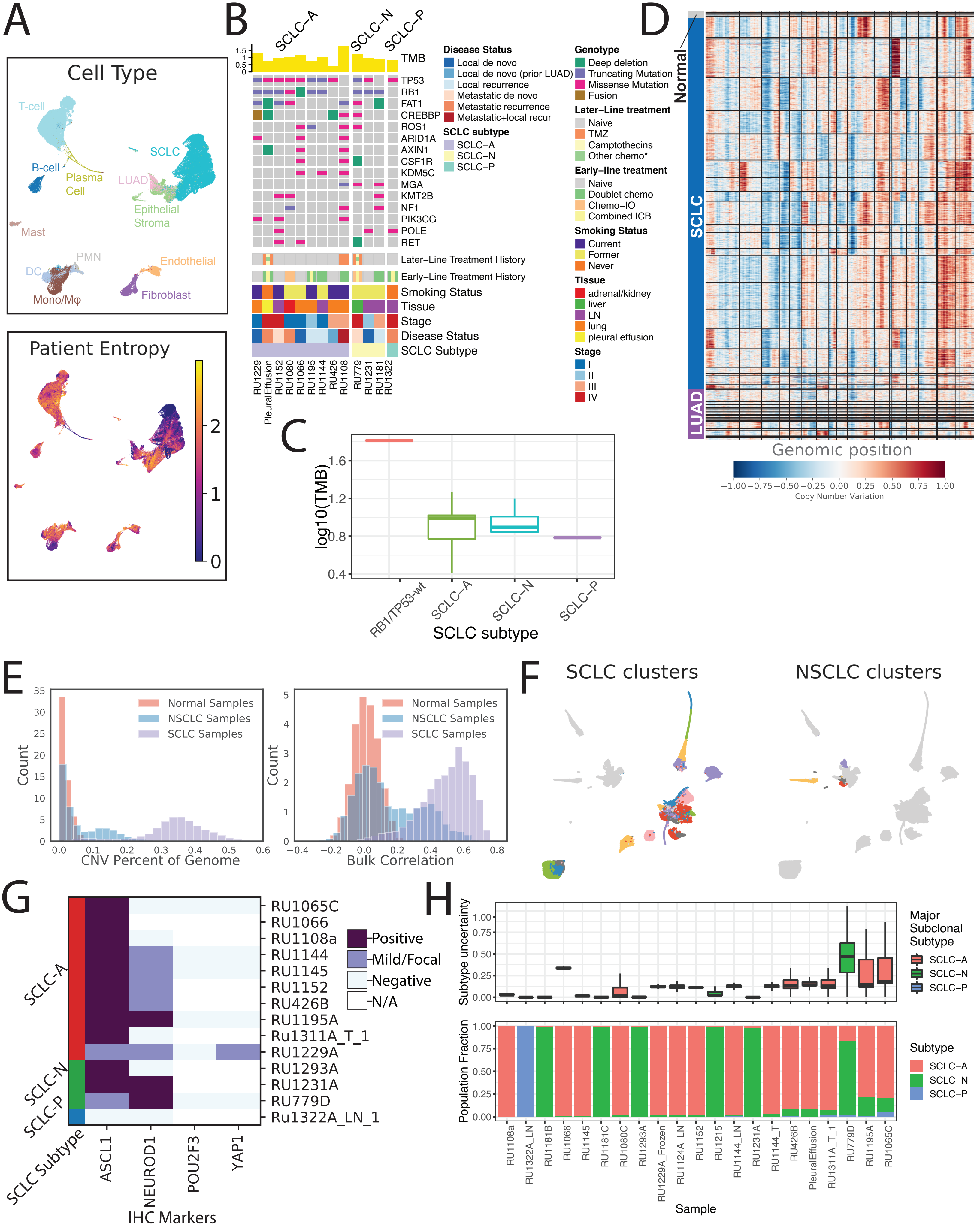

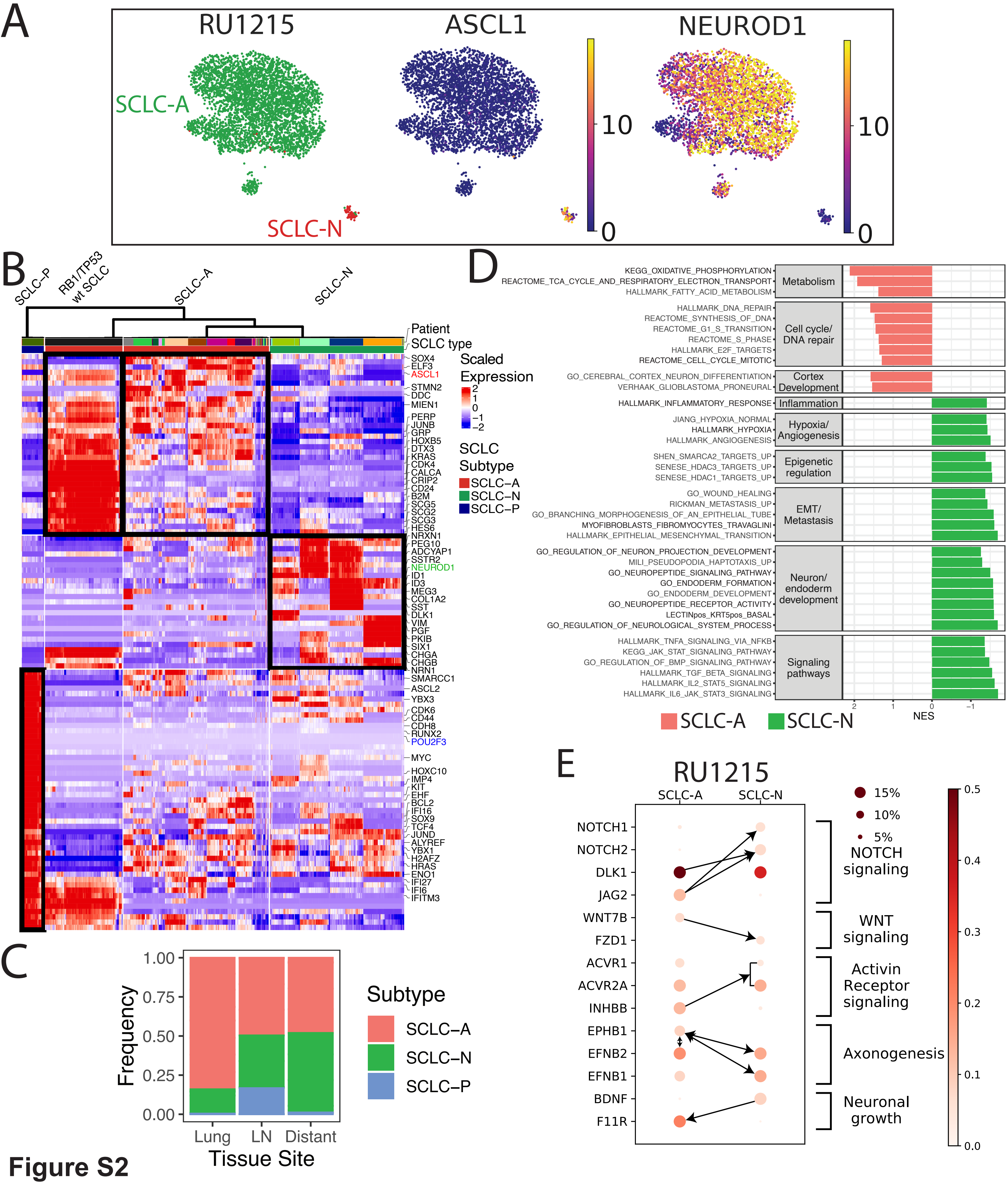

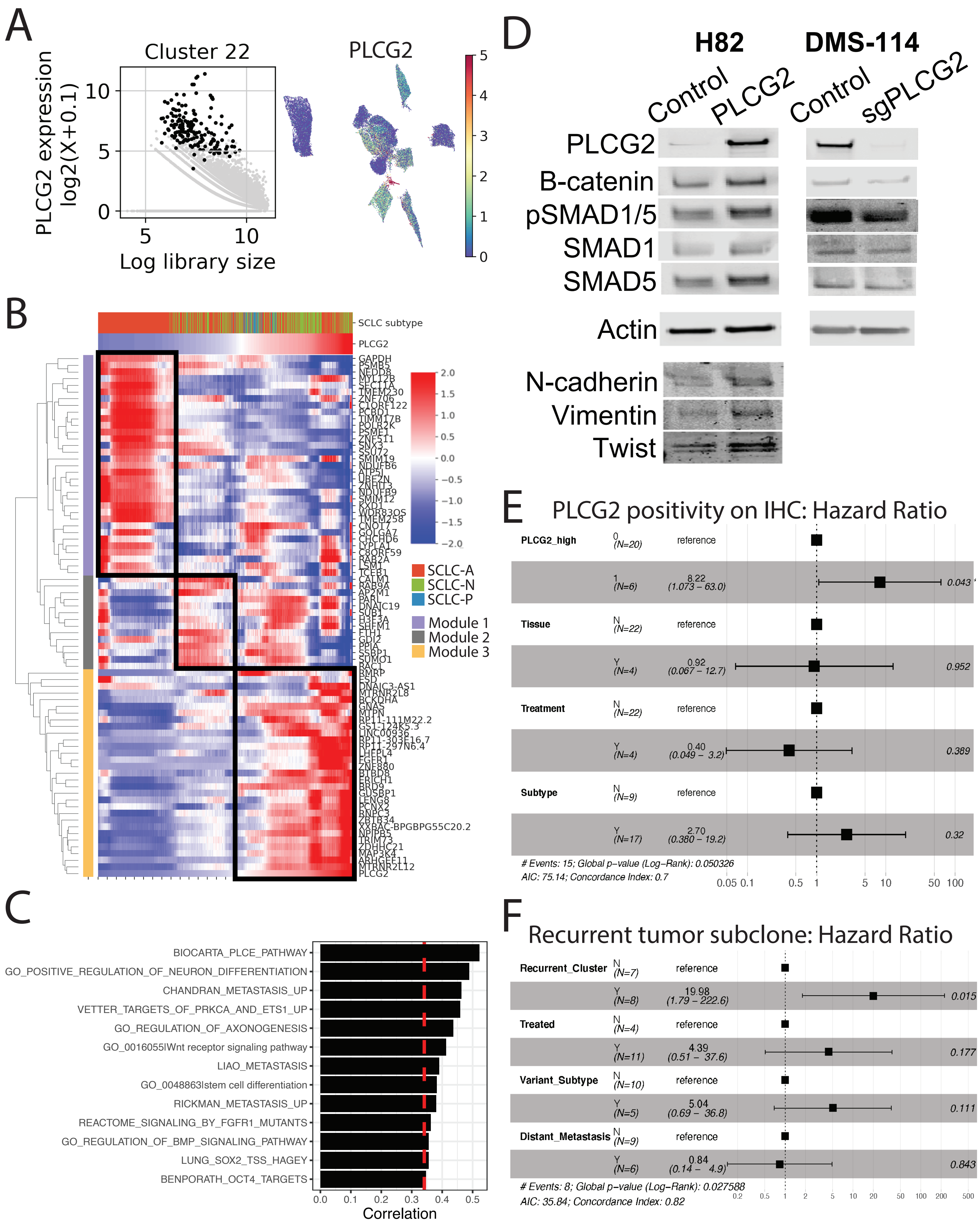

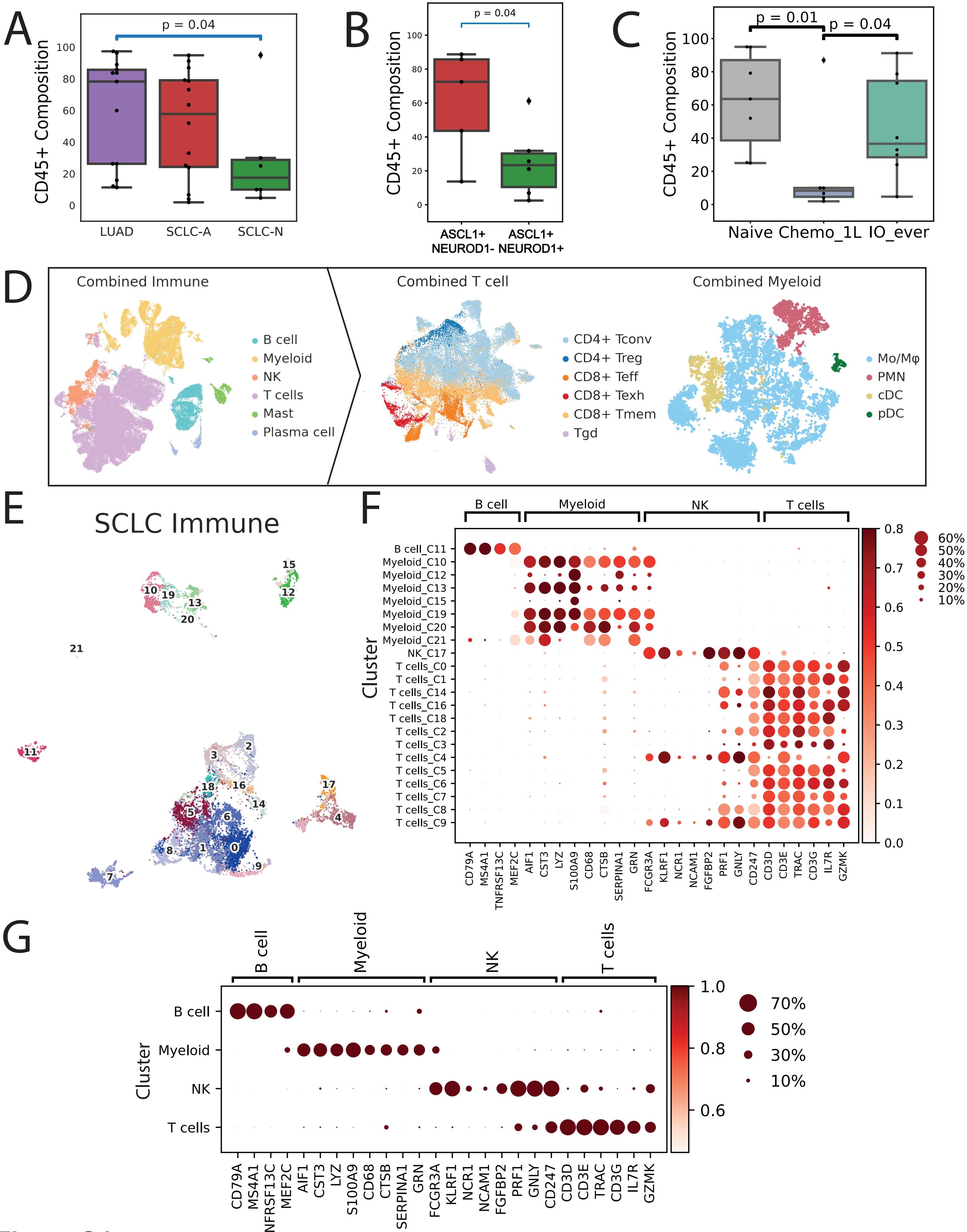

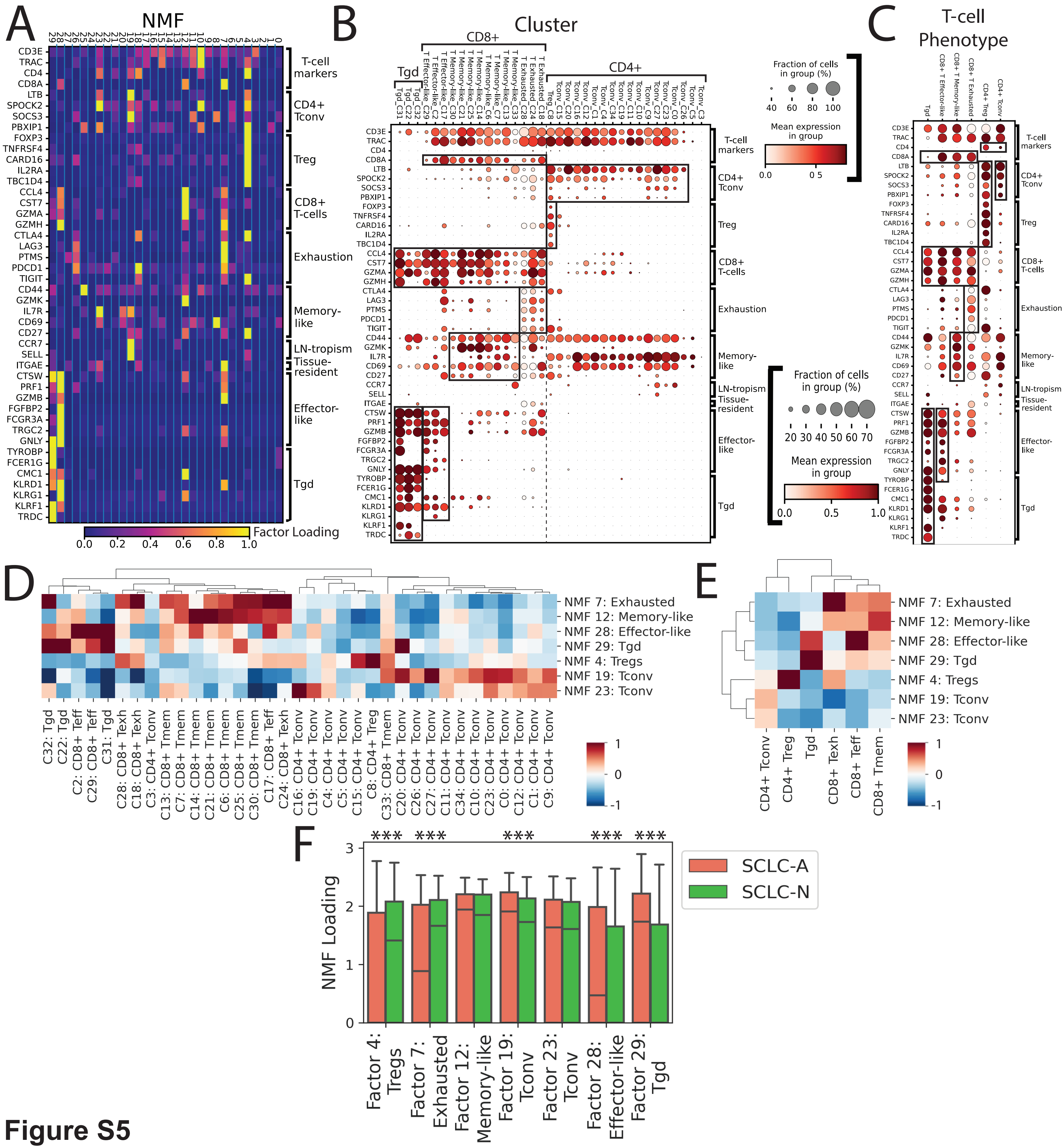

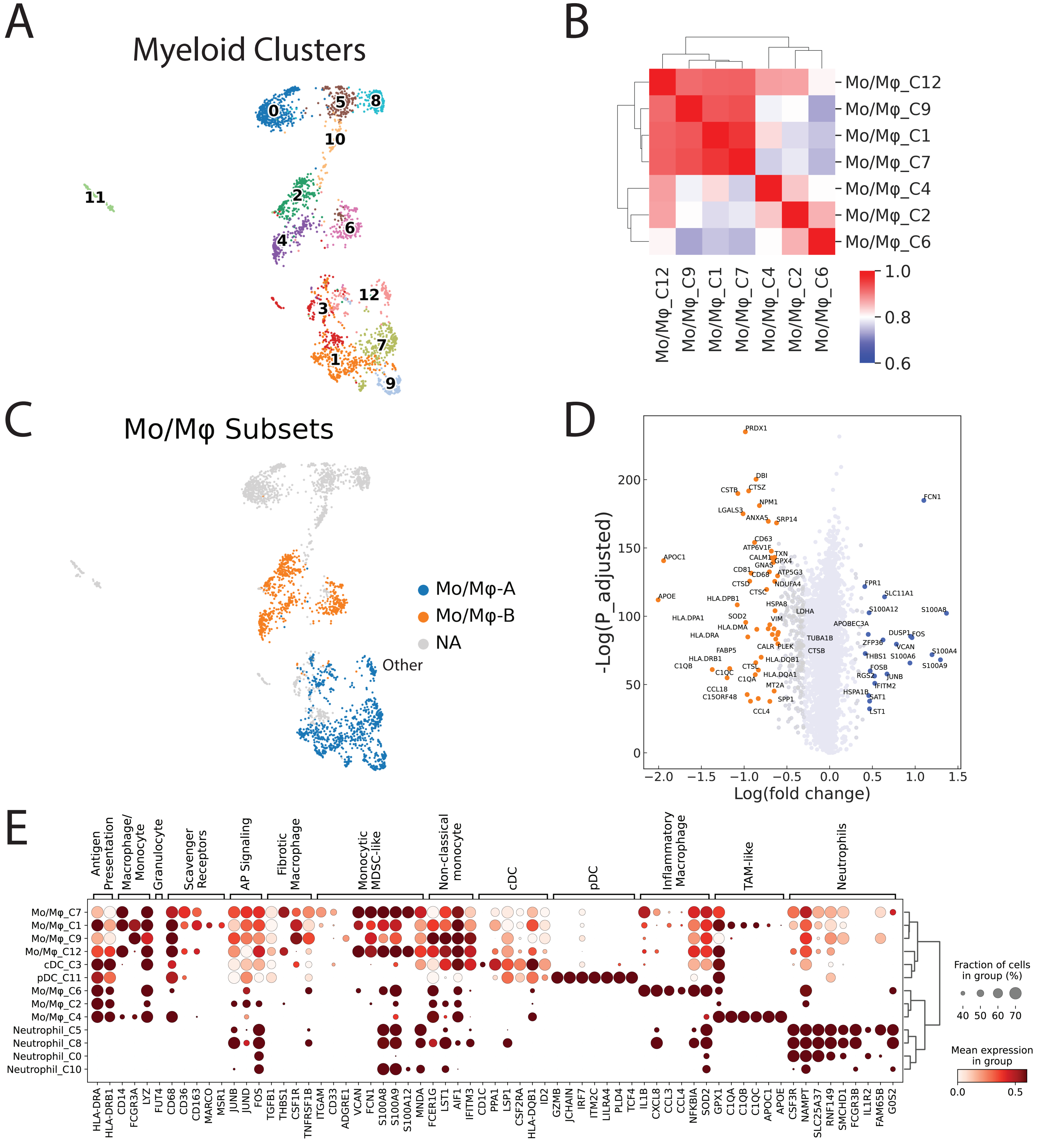

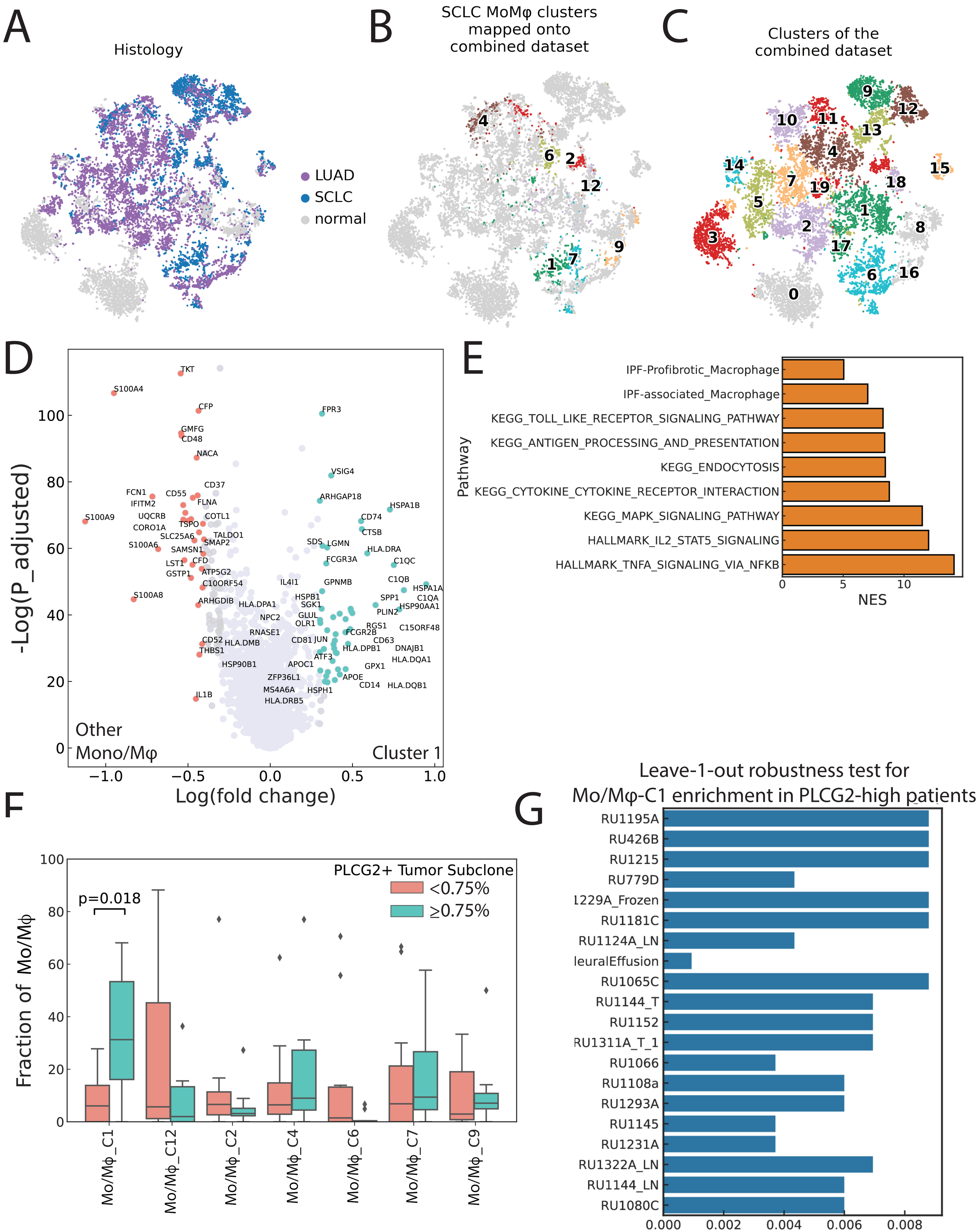
